# NK and NKT cells in the pathogenesis of Hidradenitis suppurativa: Novel therapeutic strategy through targeting of CD2

**DOI:** 10.1101/2023.10.31.565057

**Authors:** Mahendra P. Kashyap, Bharat Mishra, Rajesh Sinha, Lin Jin, Nilesh Kumar, Kayla F. Goliwas, Jessy Deshane, Boni E. Elewski, Craig A. Elmets, Mohammad Athar, M. Shahid Mukhtar, Chander Raman

## Abstract

Hidradenitis suppurativa (HS) is a chronic debilitating inflammatory skin disease with poorly understood pathogenesis. Single-cell RNAseq analysis of HS lesional and healthy individual skins revealed that NKT and NK cell populations were greatly expanded in HS, and they expressed elevated CD2, an activation receptor. Immunohistochemistry analyses confirmed significantly expanded numbers of CD2+ cells distributed throughout HS lesional tissue, and many co-expressed the NK marker, CD56. While CD4+ T cells were expanded in HS, CD8 T cells were rare. CD20+ B cells in HS were localized within tertiary follicle like structures. Immunofluorescence microscopy showed that NK cells (CD2^+^CD56^dim^) expressing perforin, granzymes A and B were enriched within the hyperplastic follicular epidermis and tunnels of HS and juxtaposed with apoptotic cells. In contrast, NKT cells (CD2^+^CD3^+^CD56^bright^) primarily expressed granzyme A and were associated with α-SMA expressing fibroblasts within the fibrotic regions of the hypodermis. Keratinocytes and fibroblasts expressed high levels of CD58 (CD2 ligand) and they interacted with CD2 expressing NKT and NK cells. The NKT/NK maturation and activating cytokines, IL-12, IL-15 and IL-18, were significantly elevated in HS. Inhibition of cognate CD2-CD58 interaction with blocking anti-CD2 mAb in HS skin organotypic cultures resulted in a profound reduction of the inflammatory gene signature and secretion of inflammatory cytokines and chemokines in the culture supernate. In summary, we show that a cellular network of heterogenous NKT and NK cell populations drives inflammation, tunnel formation and fibrosis in the pathogenesis of HS. Furthermore, CD2 blockade is a viable immunotherapeutic approach for the management of HS.

## Introduction

Hidradenitis suppurativa (HS) or acne inversa is a chronic inflammatory skin disease characterized by deep-seated lesions often in conjunction with obstruction of the hair follicles and associated hypertrophic scarring in apocrine gland-bearing regions of the body (*1–3*). Although the disease is underdiagnosed, currently, the estimated world-wide prevalence is 1-4% with a female-to-male ratio of 3:1 (*4, 5*). The mean onset of the disease is 21.8 years and individuals of African ancestry are more commonly affected than those of European ancestry (*6–9*). The clinical features of the disease include painful nodules, malodorous discharge, recurrent abscesses and development of scars and fibrosis (*1, 2, 10, 11*). The affected primary regions are the groin, axillae and anal folds. Secondary affected regions are submammary skin tissue and skin folds particularly is obese individuals. The clinical features have a dramatically negative effect on the quality of life (QOL) often contributing to depression and anxiety in the patients. Invasive squamous cell cancer (SCC) is considered one of the most severe complications of HS which carries a very high risk of death (*12*).

HS is classified by Hurley stages I, II and III, based on disease severity and extent of affected area (*6*). Although inflammation is a key feature, a major gap in knowledge includes the mechanistic underpinning of the disease onset, progression, and overall etiology. The immune cell populations implicated in HS have been T cells (CD4, CD8), B cell populations (B cells and plasma cells), neutrophils and macrophages (*10, 13–18*). CD4+ effectors were mainly Th1 and Th17 as determined by flow cytometry and gene expression profile HS lesional skin (*2, 19–22*). Consistent with this, Th1 (and CD8^+^ T cell) signature cytokines, IFN-γ and TNF-α, and Th17 signature cytokines IL-17A, IL-17F, IL-22 and IL-6 are elevated in HS (*1, 23–27*). Neutrophils in HS lesions are likely recruited by IL-17, IL-8 and/or infection (*13, 14, 28*). Neutrophils form neutrophil extracellular traps (NETs) within HS lesions promoting *in situ* immune dysregulation and amplification of the inflammatory cascade through activation of the NLRP3-inflammasome pathway and/or suppression of immune regulatory mechanisms (*13*). The inflammatory milieu of HS recruits monocytes and promote their differentiation into inflammatory M1-like macrophages (*17, 18, 29, 30*).

The most common treatment for HS is systemic antibiotics and intralesional corticosteroids to treat infection and attenuate inflammation (*1, 31*). TNF-α blockers are and anti-IL-17a (secukinumab) are biologics for the treatment of HS with benefit for treating patients with refractory disease (*15, 32, 33*). Targeting IL-17F, IL-1β and JAK for HS are being examined in clinical trials (*34–39*). However, all these treatments at best provide marginal relief. Hence there is an essential unmet need for new treatment for HS.

Our goal in this study was to obtain an understanding of the etiology of HS and its molecular underpinnings that could serve as a basis to translate the information into a highly effective therapy. We employed a multi-omics approach including single-cell and bulk transcriptomes, and regulome to build deterministic associations among multiple biological factors in HS. We discovered that NK cell populations, CD3^+^CD56^bright^ NKT cells and classical cytolytic CD56^dim^ NK cells, are major innate lymphoid cells contributing to HS disease. These NKT cells and NK cells were partitioned at different locations within HS skin contributing to different aspects of HS pathogenesis. Molecularly, the NKT and NK cells expressed high levels of the co-stimulation/adhesion molecule CD2, and its blockade attenuated the inflammatory gene and cytokine expression profile of HS.

## Results

### NKT cell populations are major pathogenic immune cells in HS skin

We performed single-cell RNA seq (scRNAseq) analysis with cells from control (n=4) and HS (lesional; n=6) skin and the dataset were log normalized, integrated with harmony and dimensionally reduced using the UMAP algorithm (Table S1; Fig. S1A)(*40, 41*). A total of 27,442 cells were grouped into 23 clusters and following unsupervised analysis, they were annotated to populations/subpopulations of adaptive and innate, keratinocytes, fibroblasts, endothelial cells, and melanocytes (Fig. 1A, Fig. S1B, Table S2) (*16, 42*). T cells, B cells and three NKT cell populations (NKT, NKT-CD8, NKT-MAIT) were the major lymphoid cell populations (Fig. 1A, Fig. S1C). The NKT-CD8 and NKT-MAIT clusters share features with CD8 T cells and MAIT cells, respectively, but a larger proportion of the cells within these clusters were closely related to NKT cells (Fig. 1A, Fig.S1C). The proportion analysis revealed a striking expansion of NKT cells and NK in HS, in addition to many other immune cell populations compared to normal skin (NS) (Fig. 1B). This feature was present in each of the six patients (Fig. S1D and Table S1). The potential importance of T cells and B cells in HS pathogenesis was previously reported (Fig.1B) (*16*).

**Figure 1:**
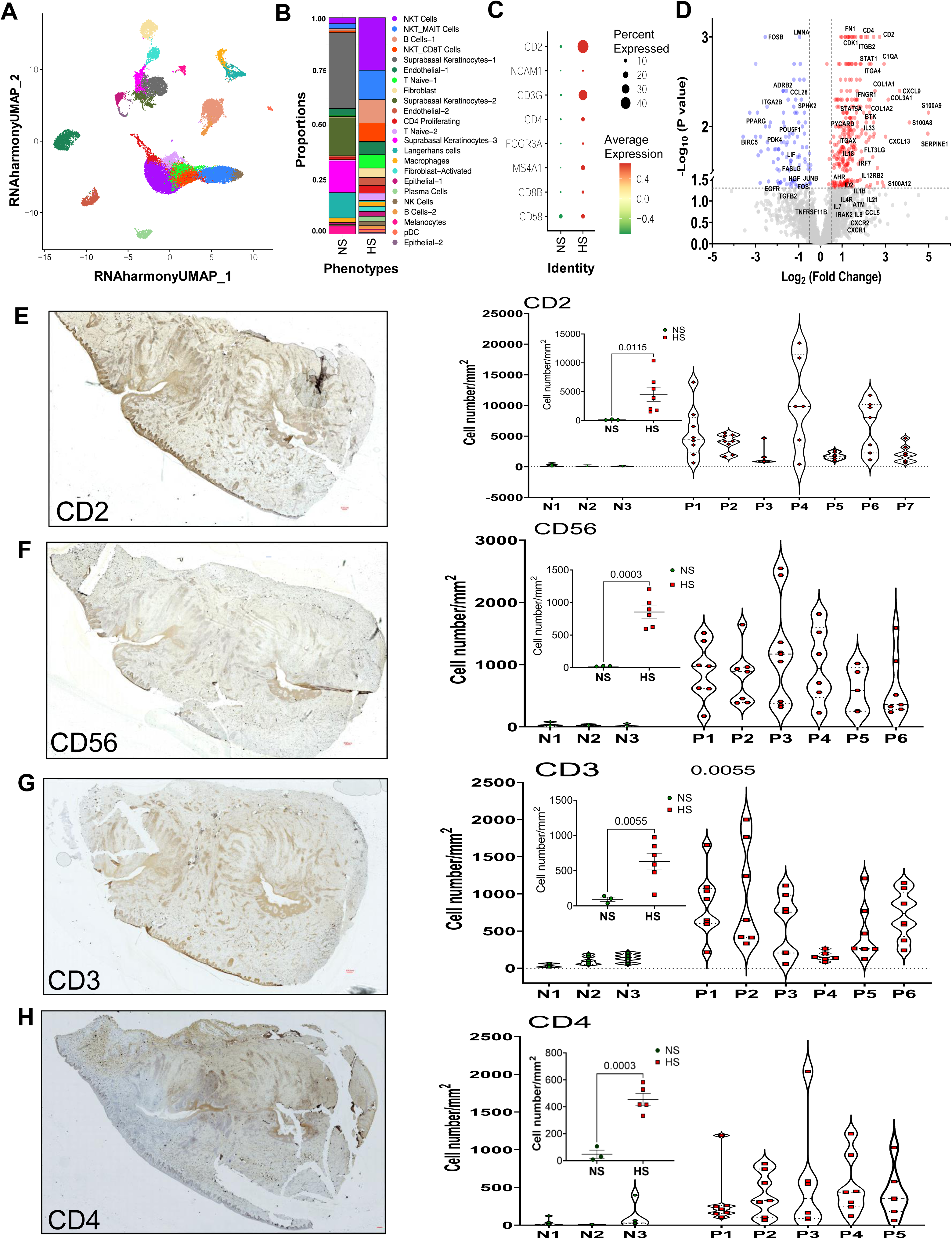
NKT and NK cell populations as major innate immune cells in HS skin. **(A)** UMAP of scRNAseq transcriptomes from skin of normal skin (NS; n=4) and HS (n=6). Seurat 4.0 was used to identify 23 clusters representing immune and non-immune cell populations. **(B)** Proportion of annotated cell type in each cluster from NS and HS. **(C)** Dot plot depicting normalized gene expression in NS and HS for a panel of genes. **(D)** Volcano plots of differentially expressed genes in HS (n=6-8) compared to NS (n=8). **(E-H)** IHC image analysis of NS and HS skin samples showing absolute number of CD2 cells, CD56 cells, CD3 cells and CD4 T cells. The images (left micrographs) are representative from HS. Red bar in each micrograph is 300 μM. NS representative micrographs are shown in Fig. S4A. Each dot in the violin plots represent number of cells/region representing ≤400 20X images from NS or HS as indicated. The inset graph within each violin plot is mean±SEM for NS or HS, each dot represents the average from the violin plots. For a comprehensive analysis the entire area of HS lesional skin and NS as shown in the micrograph was scanned and between 2000 and 4000 images were obtained for each sample. The images were stitched to generate single composite image (shown) and immune cells were counted using Macro Cell Count module of BZ-X analyze (Keyence Corporation) which count several hundred images simultaneously in single run. Based on tissue size each stitched image was split in 3-8 parts with each part containing ≤ 400 images and counted using the same software settings. Area of each image was also measured and cell number is extrapolated as number of cells present per millimeter square area.

The expression levels of several genes associated with lineage and/or function were expressed within HS immune cell populations (Fig. 1C, Fig. S1C, Table S2). Among them, the elevated expression of CD2, a cell surface receptor associated with NKT, NK and T cell activation (*43–47*) was highly expressed on 40% of cells from HS (Fig.1C). The expression levels of NCAM1 (CD56), CD3G (CD3γ), CD4, MS4A1 (CD20), CD8B (CD8β) and CD58 (LFA3) were also elevated in HS over controls. The monocyte/macrophage cell cluster exhibited an inflammatory gene signature with elevated expression of STAT1, IFNGR1, IFNGR2, IRF7, genes associated with activation of type 1 and type II IFN signaling (Table S2). This cluster and that of keratinocytes had enhanced expression of IL18, a cytokine that promotes T cell, NKT cell and NK cell activation and expression of IFN-γ (*48–52*).

Next, we compared our scRNAseq data findings to bulk transcriptome profile from four independent public data sets, GSE151243, GSE154773, GSE79150, and GSE128637. The analysis of differentially expressed genes (DEGs) found that a total 1,614, 2,080, 434, and 359 genes were significantly upregulated, and 1,354, 1,747, 309, and 300, were significantly downregulated in HS, respectively (Fig. S2A). KEGG analysis revealed that most of the upregulated DEGs involved immune response, cell activation, stress-response, inflammatory response, and IL 17 signaling. Whereas most of the downregulated DEGs represented cell differentiation, lipid metabolism and ion transport. Relevant to this study, using Ingenuity Pathway Analysis (IPA), we noted that NK cell signaling, among others, was a significantly activated pathway in HS (Fig. S2B).

To validate the scRNA data and to further interrogate the differentially expressed genes between HS and controls, we performed a qRT-PCR gene expression array using skin tissue from a cohort of eight HS and six-eight healthy controls (Tables S3, S4). Of the eight HS tissues analyzed in this assay, six were the same individuals used for scRNAseq (Fig. S3A). The qRT-PCR OpenArrays representing 2429 gene targets, showed that 170 genes were significantly (Log2FC≥1; P<0.05) elevated and 66 genes were significantly (Log2FC ≤ − 1; P<0.05) down modulated in HS compared to controls (Fig.1D; Fig. S3B, C and Tables S3, S4). CD2, S100 gene family, kinase BTK, integrin ITGAX, complement protein C1QA and several cytokines/chemokines and their receptors, phosphatases, kinases, and other molecules associated with an inflammatory signature were among the genes whose expression was significantly elevated in HS compared to controls (Fig.1D; Tables S3, S4). We recently reported the discovery of a keratinocyte population specific to HS characterized by enhanced expression of S100 gene family driving expression of inflammatory genes (*53*).

### Elevated CD2 expression and distinct distribution of NKT cells and NK cells in HS lesions

The finding based on scRNAseq analysis shows that NKT cell populations are greatly expanded in HS and they along with CD4 T cells express high levels of the activation molecule CD2 is novel (Figs.1A, B, C, Table S2). To validate this finding, we rigorously analyzed serial sections of skin from HS patients (n=5-7) and controls (n=3) stained for CD2, CD56, CD3 and CD4 expressing cells by immunohistochemistry (IHC) (Figs. 1E-H). Large numbers of cells expressing high levels of CD2 were distributed across the entire HS section (Fig. 1E). This feature was observed in each of the seven HS sections analyzed and total numbers of CD2 expressing cells were exponentially higher in HS than that in NS (Fig. 1E, Fig. S4A). Similarly, the numbers of CD56, CD3 and CD4 expressing cells were significantly greater in HS vs NS (Figures 1F-H). NKT cells are CD56^bright^ and CD3^+^ and classical mature (highly cytolytic) NK cells are CD56^dim^ (*54*). The CD56 staining in IHC in these HS skin sections primarily represent CD5^bright^ NKT cells and hence most co-express CD3 as seen in the representative serial section (Fig.1F, G). The IHC staining is not sensitive enough to accurately detect CD56^dim^ cells. As expected, HS sections contained significantly greater numbers of CD4^+^ T cells than controls, but they were fewer than CD56 expressing cells (Fig.1H).

NKT cells and NK cells are heterogenous and have functions determined by tissue and/or organ localization (*54, 55*). We, therefore, interrogated by immunofluorescent microscopy if their spatial localization in HS skin corresponds to distinct functions in HS pathogenesis. H&E stained serial sections from normal and HS skin were used to localize cells expressing CD56 (NKT, NK), CD3 (T and NKT) and CD2 (T, NKT and NK) by immunofluorescence microscopy. The tissue morphology of skin from healthy controls revealed well-defined epidermal and dermal regions, in contrast, HS skin (Hurley late stage II and stage III) showed highly inflamed areas with hyperproliferation of the epidermal and dermal compartments (Fig. 2A, Fig. S4B). Histology showed the presence of tunnels protruding deep inside the dermis and extending to the hypodermis with extensive cellular infiltration of different populations of immune cells. NS contained very few T cells (CD3^+^CD56^-^) and NKT (CD3^+^CD56^bright^) or classical NK cells (CD56^dim^) (Fig.2A, B). In contrast, HS skin contained large numbers of classical NK (CD56^dim^), NKT (CD3^+^CD56^bright^) and T cells (CD3^+^CD56^-^). CD2 was highly expressed on all NK, NKT and T cells. Classical NK cells (CD3^-^CD56^dim^) were predominantly present within the epidermal and dermal regions of HS skin (Fig. 2A) whereas NKT (CD3^+^CD56^bright^) cells were enriched adjacent to the tunnels and hair follicular regions (Fig.2A). Quantification shows that the epidermal and dermal regions of HS contain significantly less NKT (CD3^+^CD56^bright^) cells than tunnel areas (Fig. 2B). To further distinguish between T cells and NK populations, we stained for expression of CD16 (FcγRIII), a marker that is expressed on NKT and NK cells but not T cells (*54*). We observe that most cells in HS express CD16 along with CD56 establishing that these were indeed NK cell populations. Epithelial cells in HS expressed greatly elevated expression of CD58 (LFA3), the ligand of CD2, compared to normal skin (Fig. 2C) (*44, 56, 57*). CD2 expressing cells interacted with CD58 expressing epithelial cells. Our data shows that NKT cells and NK cells are major lymphocyte populations within HS skin, and they are spatially localized in distinct regions of HS lesions suggesting independent roles in features of disease pathogenesis.

**Figure 2:**
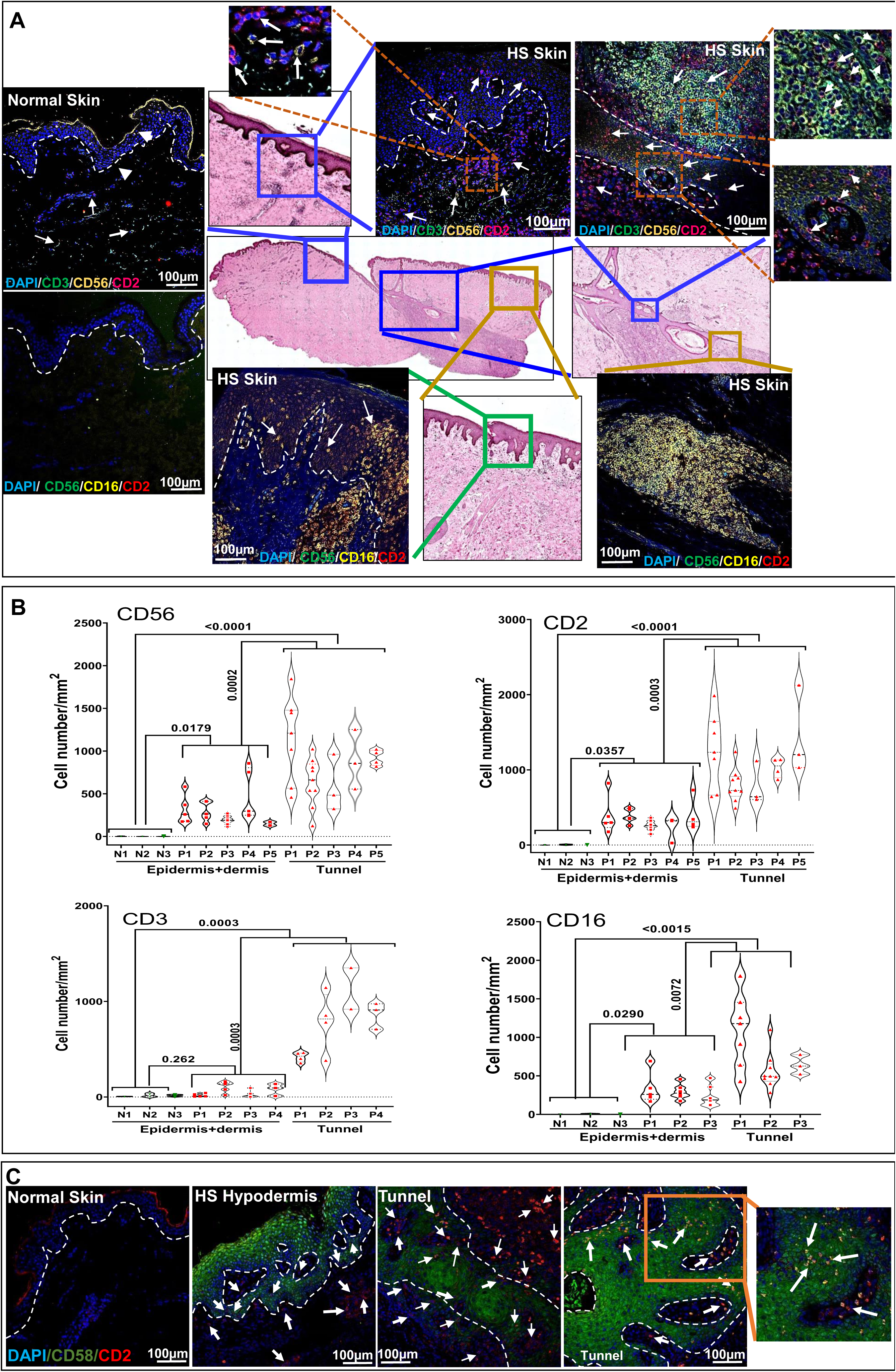
NKT and NK cells are spatially localized in different regions of HS Skin. (**A**) Immunofluorescence staining for CD3, CD56 and CD2 (upper panel) and, CD2, CD3 and CD16 expressing cells (lower panel) in normal and HS - skin as indicated. H&E staining shows “stitched skin” (**center**) with boxes denoting magnified areas of epidermis/hypodermis and sinus tract in HS. Scale bar of immunofluorescence images is 100 µm. Arrows in the immunofluorescence images point to CD2^+^CD3^+^CD56^bright^ (NKT cells) and CD2^+^CD3^-^CD56^dim^ (mature NK cells) in the epidermal/hypodermal and Tunnel regions. T cells in the immunofluorescence images are CD2^+^CD3^bright^CD56^-^. Tunnels in HS show cluster of NKT cells that is spatially separated from NK cells. (**B**) Graph showing absolute number of CD56, CD2, CD3 and CD16 cells in upper epidermal/dermal region and near tunnel regions. CellSense software was used for absolute cell counting. Minimum 20 confocal images/individual (20x) were counted comprehensive analysis without biased selection of regions. (**C**) Lower panel show CD2 cells interacting with CD58 (keratinocyte) expressing cells. Data is representative of HS skin (n=3) and normal skin (n=3).

### B lineage and CD8 T cells in HS skin

Several recent studies have shown that B cells are present in elevated numbers within HS lesions and likely contribute to pathogenesis of disease (*58*). We analyzed by IHC for CD20^+^ B cells in HS (n=6) and control (n=3) skin tissue, representing serial sections from the same individuals as in Fig. 1E. In HS B cells were localized to tertiary follicle like structures and were significantly greater than that in controls (Fig. 3A). However, they were a lot fewer than CD56 expressing cells (Fig. 3A vs Fig. 1F). Immunofluorescence staining for B cells (CD20^+^) and plasma cells (IgG^+^) showed that they were much fewer in number than CD2^+^ cells in HS, thereby recapitulating the IHC data (Fig. 3B vs 2B, Fig. S5).

**Figure 3:**
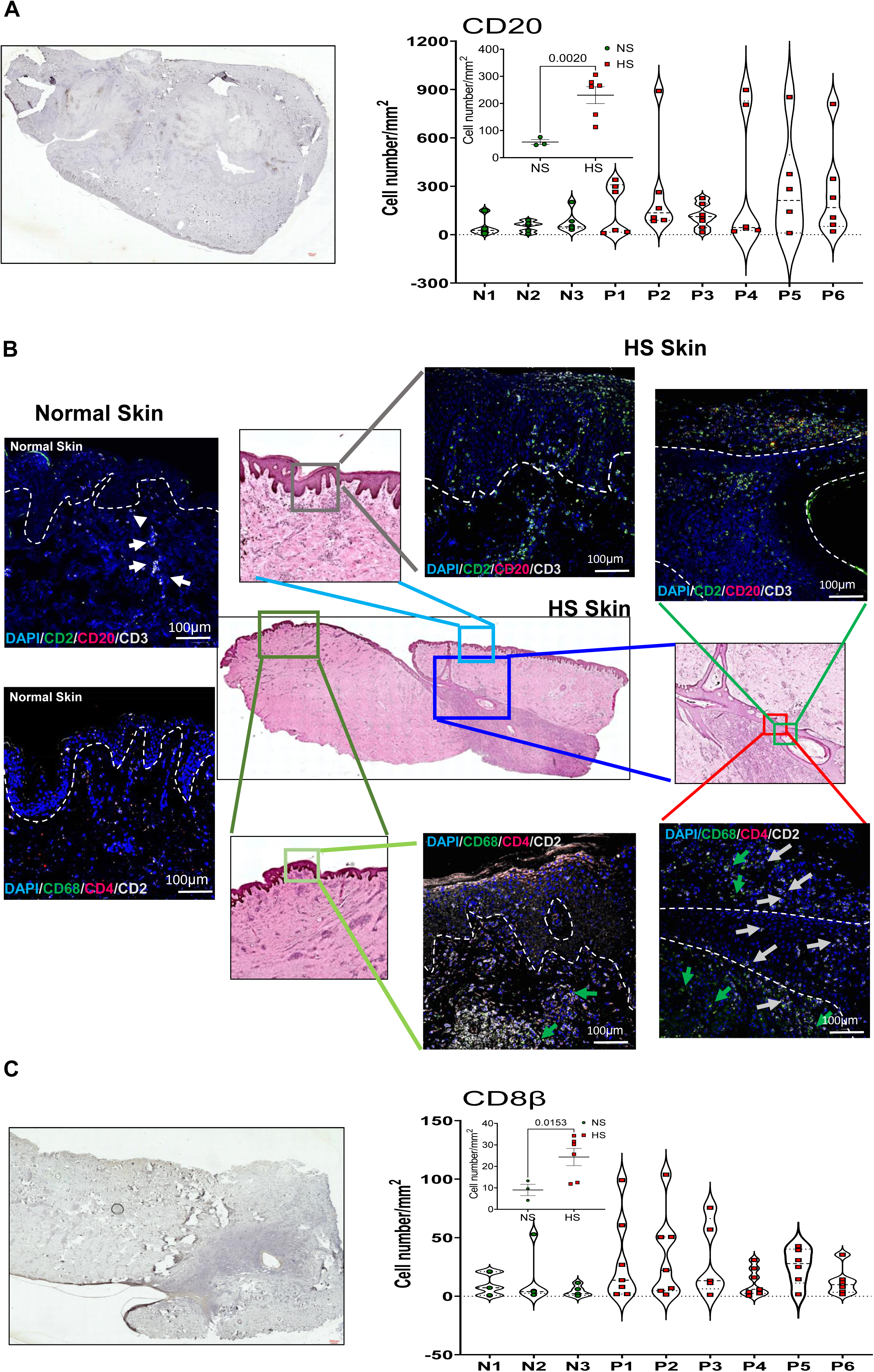
B cells and CD8 T cells in HS skin. (**A**) IHC image analysis of normal (n=3) and HS skin (n=6) samples showing absolute number of CD20+ B cells. Cell number was determined as in Fig 1. (**B**) Immunofluorescence micrographs of skin stained with (upper panels) antibodies to CD20, CD2 and CD3; (lower panels) antibodies to CD4, CD68 and CD2. (**C**) IHC image analysis for CD8β expressing cells. Representative IHC micrograph shown. Violin plot graph NS (n=3) and HS skin (n=6) and inset graph as described in Fig. 1.

The T cells in epidermal, dermal and tunnels of HS skin were predominantly CD4^+^; CD8^+^ T cells were infrequent (Fig. 3B, Fig. S5). Skin sections stained for CD8β by IHC showed that although HS had significantly greater numbers of CD8 T cells, the overall numbers were very small (<40 cells/mm^2^) (Fig. 3C). This is consistent with other reports (*15, 16*). As previously reported, HS skin also contained an expanded proportion of macrophages (CD68^+^, EMR1^+^) and neutrophils (MPO^+^) (Fig. 3B, Fig. S6) (*13, 17, 18, 29*). MPO^+^ cells were mostly localized in hypodermal regions and areas of abscess formation.

### Distinct roles of NK and NKT populations in HS pathogenesis

Following the demonstration that NKT and NK cell populations are spatially localized in different regions, we predicted that they may have distinct roles in HS pathogenesis. Perforin and granzymes (granzyme A and granzyme B) are proteins necessary for NKT cell and NK cell mediated cytotoxic activity of target cells (*54, 59, 60*). Normal skin contained a small frequency of perforin-1, granzyme A and granzyme B expressing cells (Fig. 4, A-C). In contrast, HS skin had large numbers of perforin-1, granzyme A and granzyme B expressing classical NK cells (CD3^-^CD56^dim^) within epidermis and dermis and along the border region of tunnels that seemed to be linked to their development (Fig. 4, A-C). This region of HS tissue, are enriched in TUNEL+ cells (detailed below), showing a mechanistic underpinning for tunnel formation. NKT cells and NK cells expressed equivalent levels of granzyme A, however, expression of perforin-1 and granzyme B was distinctly lower in NKT cells than NK cells. The immunofluorescence staining images reaffirm that NKT cells and NK cells in HS express high levels of CD2. A significant proportion of the CD56^dim^CD2^hi^ granzyme B^hi^ cells also expressed high levels of CD11b (MAC1/ITGAM/CR3) (Fig.4C). Overall, the differential expression of these proteins in NK populations from different regions of HS skin reveals a heterogeneity in NKT and NK cell populations. We extracted the scRNAseq data of immune cells clusters (NKT Cells, NKT_MAIT Cells, NKT_CD8T Cells, T Naive-1, CD4 Proliferating, T Naive-2, and NK Cells) (Fig. 1A) and performed subset analysis. Using the re-clustering and unsupervised annotation, we identified 10 cell populations representing six cytolytic cell populations: two NKT, two CD8 (CD3+/^-^), one NK and one with MAIT characteristics (CD3^+^CD161^+^) (Fig. 4D, 4E). This affirms the existence of heterogenous NKT and NK cell populations in HS. Taken together these findings indicate the different roles of NKT and NK cells in HS pathogenesis.

**Figure 4:**
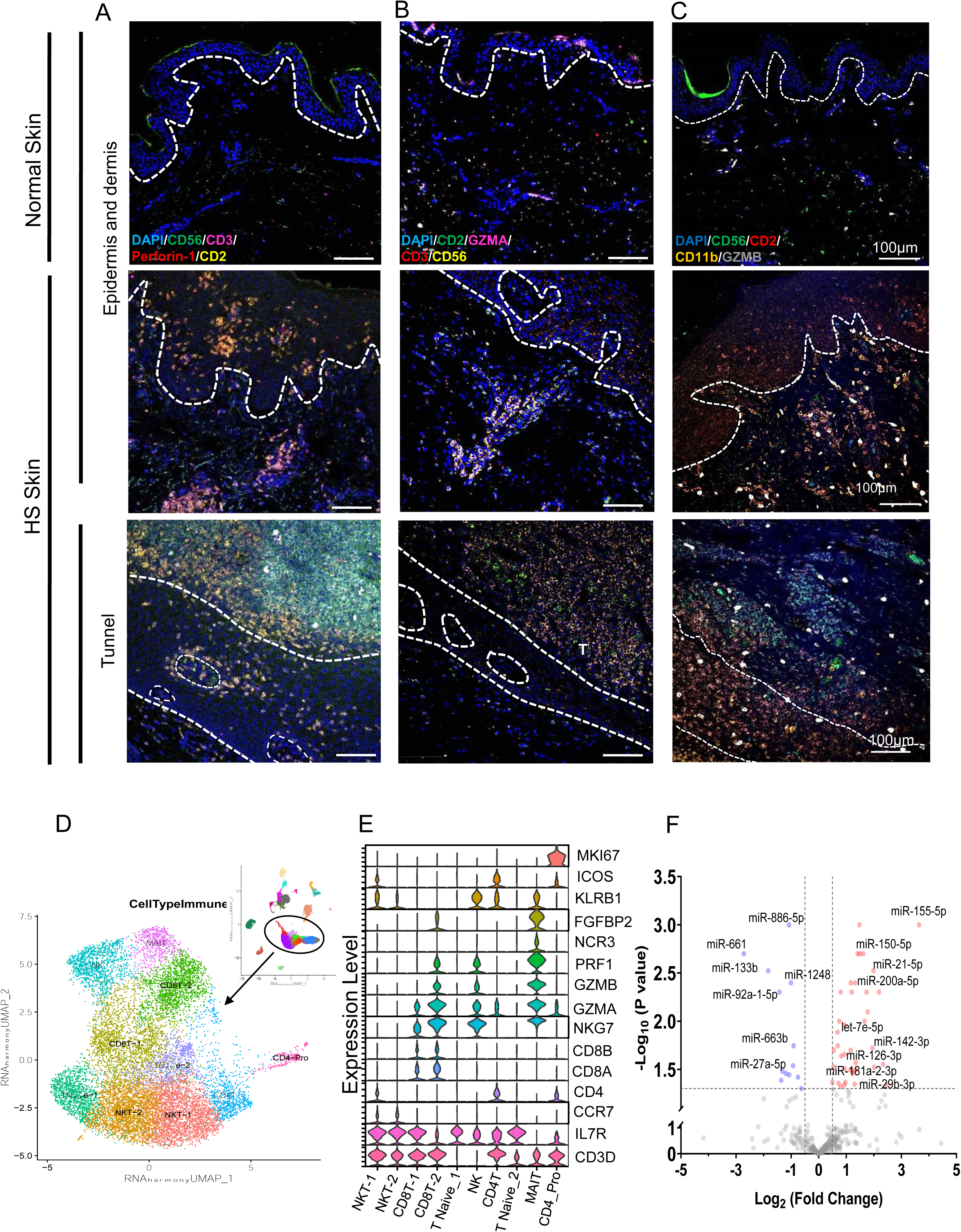
Heterogenous NKT and NK cell populations defined by expression of perforin, granzyme A, granzyme B and CD11b. (**A**) Perforin expressing NKT (CD3^+^ CD2^+^CD56^bright^) and NK (CD2^+^CD56^dim^) cells in normal skin and HS skin. Perforin expressing NKT and NK cells are prominent in the epidermis, hypodermis, tunnel and border regions of tunnel. NK cells (CD3^-^) express higher levels of perforin NK2 cells (CD3^+^). (**B**) Granzyme A expressing NKT and NK cells in normal skin and HS skin. Levels of granzyme A are equivalent in NKT and NK cells in HS. (**C**) CD11b and granzyme B expressing cells in normal skin and HS skin. Some of the NK cells (CD56^dim^), but not NKT cells in HS express CD11b. Granzyme B expression is observed predominantly within NK cells. (**D**) UMAP of 10 immune cell sub-clusters extracted from scRNASeq analysis in Figure 1A. (**E**) Expression of NKT and NK associated genes in each sub-cluster. (**F**) **Volcano plot of expressed miRNAs in normal and HS skin.** miRNA profile in skin from normal (n=6) and HS (n=9) were determined using OpenArray panel containing 754 well characterized miRs. 27 miRs were significantly upregulated while 7 miRs were significantly downregulated. The eleven miRNAs with known function in modulating NK cell differentiation and/or function is labeled in the volcano plot.

### MicroRNA profile reflects NK cell differentiation and/or activation

To interrogate the transcriptional regulome, we performed global microRNAs (miRs) analysis with tissue from nine HS and six healthy controls (Fig. 4F and Fig. S7A, Table S5. S6). We identified 36 differentially expressed miRs in HS compared to healthy skin of which 27 miRs were significantly upregulated and 9 miRs were significantly down-regulated (log_2_FC > |1|, *p*-value<0.05;) (Fig. 4F, and Table S6). Eleven (11) of the 36 differentially expressed miRs (mIR-150, miR-27a5p, miR-155, miR-21, miR-142-3P, miR-126, miR-29b, Let7, and miR-200a) have a role in NK cell differentiation and/or function (Figs. S7B, 7C) (*61, 62*). Particularly, miRNAs (mir-150, miR-155, miR-21, miR-200a, miR29a-5p, Let7) that enhance NKT or NK cell differentiation and/or activity were elevated in expression and miRNAs that regulate NK activity (miR-27a-5p, miR-181a-2-3p) were decreased in expression in HS relative to NS (Fig. 4F and Tables S5, S6) (*62*).

### Functionally distinct roles for NK and NKT in HS pathogenesis

Consistently, TUNEL+ cells were in the deep dermis along the peripheral regions and deep tunnels adjacent to NK cells (Fig. 5A). CD3^+^CD56^bright^ NKT cells were distinctly localized separate from NK (CD56^dim^) cells and predominantly within the outer region of tunnels. CD2^+^ lymphocytes were located primarily within the peripheral regions of the tunnels, and they expressed PAR2, a protein that counteracts apoptosis (*63, 64*). None of the apoptotic cells expressed CD2, suggesting that the NK and NKT cells may upregulate PAR2 as a mechanism to prolong survival as they cause the complex pathogenic events in HS.

**Figure 5:**
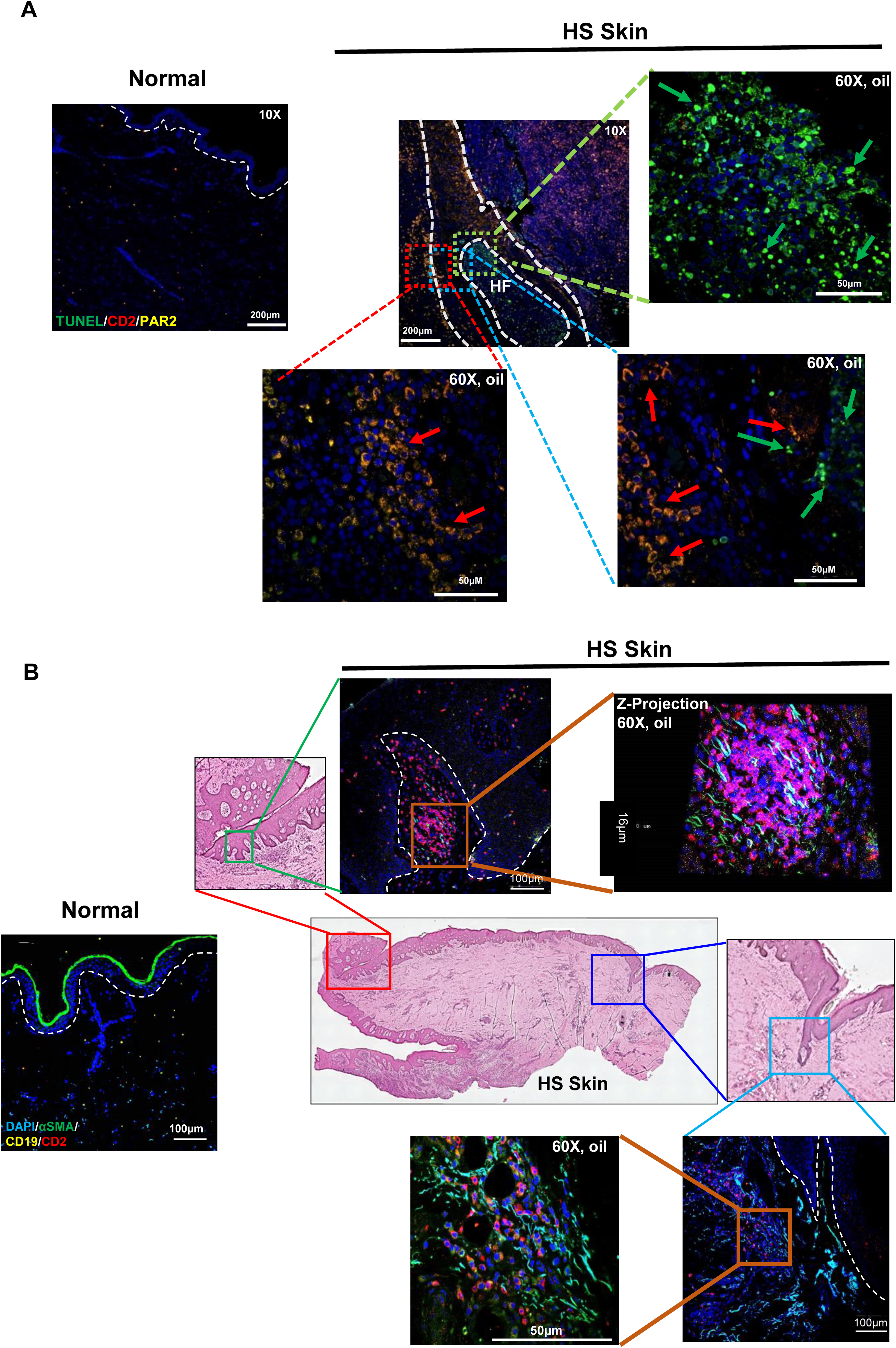
NKT and NK cells associated with pathological processes in HS. (**A**) Localization of NKT/NK cell populations (CD2^+^) in regions of ongoing apoptosis (TUNEL+). Immunofluorescence staining shows PAR2 expressing CD2^+^ lymphoid cells adjacent to cells undergoing apoptosis (TUNEL+) within sinus tracts/modified hir follicles of HS. (**B**) NKT cells interact with fibroblasts involved in ongoing fibrosis in HS. Z projected micrographs (4096X4096; 60x oil) show CD2 expressing NKT cells physically interact with αSMA expressing fibroblast like cells within modified hair follicles (upper panels). Very few CD19^+^ cells are seen in regions of ongoing fibrosis.

A key feature of HS disease progression is tissue remodeling and associated fibrosis (*16, 27*). Consistent with this, we observed that the expression of genes associated extracellular matrix (ECM) remodeling and dermal fibrosis, collagen type 1 (COL1A1, COL1A2), collagen type 3 (COL3A1), fibronectin (FN1), matrix metalloproteinase 9 (MMP9), secreted Frizzled-related protein 2 (SFRP2) and CXCL12 were significantly increased in HS (Tables S3, S4) (*65*). The elevated expression of alpha smooth muscle actin (αSMA) is a critical marker of fibrosis and ongoing tissue remodeling (fibroblast enriched regions) (*66, 67*). Enhanced expression of vimentin intermediate filaments in fibroblasts is a consequence of inflammatory insults, and necessary for fibrosis (*67*). We used these features of fibrosis to further investigate if NK cell populations were involved in the process. In HS we found expanded numbers of NK cells (CD2^+^) interacting with αSMA+ cells in the hypodermal area (the direct interaction is clearly demonstrated in z-projection images (Fig. 5B, movie S1). Subpopulations of NKT cells are known to play a significant role in liver fibrosis (*68, 69*). We found very few B cells (CD19^+^) associated with or in proximity of αSMA^+^ cells indicating that B cells are not directly involved in HS fibrosis (Fig. 5B). This is a distinct feature of HS fibrosis and tissue remodeling from that of systemic sclerosis, which involves B cells and plasma cells in its pathogenesis (*70*).

### NK T and NK cell recruitment and activation cytokine and chemokine expression in HS tissue

We assayed for the expression of cytokines, chemokines, and growth factors in skin tissue from controls and HS. The cytokines, IFN-γ, IL-1β, IL-4, IL-6, IL-8, IL-12p70, IL-15, IL-17a, IL-18, IL-22, IL-27, and TNF-α were significantly elevated in HS compared to controls (Figs. 6A, 6B and Fig. S8). IFN-γ along with TNF-α are the major effector cytokines expressed by NK cells and NKT cells. Keratinocytes are primary producers of IL-15 and IL-18 which were highly elevated in HS skin. These cytokines enhance NK cell cytotoxic activity as well as induce the expression of CD2 and thus play a role in NKT cells and classical NK cell mediated HS pathogenesis (*48, 51, 52, 71*). IL-12 was elevated in HS; signaling from this cytokine promotes survival and expansion of NK cells independent of IL-15 and IL-18 (*72*). The levels of IFN-α, IL-1α, IL-2, IL-7, IL-9, IL-31, and TNF-β were similar in HS and control skin (Fig. S8). IL-13 was elevated in HS skin, but its levels were not statistically different from that of the NS. Previous reports showed increased levels of Th17 cytokines in HS serum (*1, 16, 23*). However, we observed tissue expression of IL-17a, IL-22 and IL-6 were only marginally elevated in HS (Fig. 6B and Fig. S8). Several chemokines and growth factors were elevated in HS skin vs controls. These included CCL2 (MCP1), CCL3 (MIP-1α), CCL4, (MIP-1β), CCL5 (RANTES), CCL11 (Eotaxin), CXCL1 (GROα/KC), CXCL9, and CXCL10 (IP10), CXCL12 (SDF-1), NGFβ, EGF, FGF-2, HGF, and PIGF1 (Figs. 6A, 6B). The IL-1β induced cytokine/chemokine produced by keratinocytes, IL-8 (CXCL8), that plays an important function in recruiting neutrophils was significantly enhanced in HS (*73*). The growth factors BDNF, PDGF-BB, PIGF-1, VEGF and SCF, were present in HS skin but they were not significantly changed from controls (Fig. S8).

**Figure 6:**
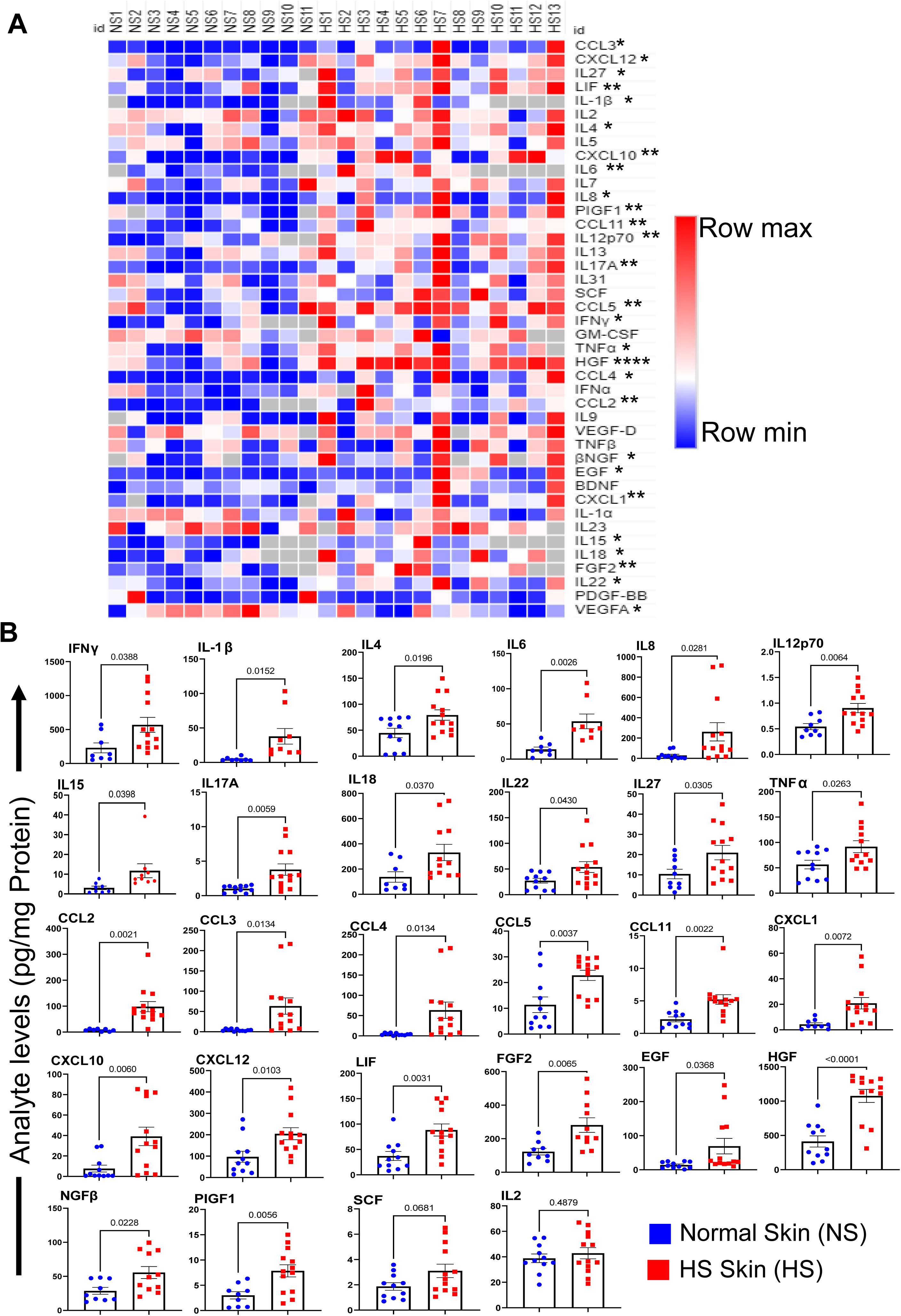
Expression of cytokines, chemokines and growth factors involved in inflammation, recruitment of effector immune cells, and cell proliferation in HS. **(A)** Heatmap showing levels of various cytokines/chemokines/growth factors in normal (n=11) and HS (n=13) skin samples. **(B)** Bar graphs with individual values of cytokine, chemokines and growth factors in NS and HS. Two tailed Student’s t-Test of normal vs HS (*P≤0.05, ** P≤0.01; ***P≤0.001; ****P≤0.0001).

### CD2 blockade in HS skin attenuates the production of cytokines, chemokines and pathogenic gene signature when tested in organotypic cultures

The data from preceding experiments suggest that distinct NKT and NK cell populations mediate type 1 (inflammatory/cytolytic) and type 2 (fibrogenic) immune activities to promote the pathogenic landscape of HS. An intricate interplay of cytokine, chemokines and growth factors produced by NK cell populations, keratinocytes and HS cell populations perpetuate the pathogenic machinery of the disease. Our results show that the upregulation of CD2 on all NK cell populations and T cells is a key feature of HS. We therefore predicted that CD2 blockade in HS is likely to reverse the expression of disease associated cytokines, chemokines, and growth factors as well as the gene expression signature.

We tested this hypothesis employing our recently developed HS skin organotypic cultures (*74*) with two experimental paradigms, CD2 blockade by itself and following challenge with LPS. We found that cultures of HS skin (n=3) treated with anti-CD2 mAb vs IgG (control) contained significantly reduced levels of cytokines, chemokines and growth factors associated with inflammatory pathogenesis that including those responsible for NK cell activation and/or maturation (Fig. 7A, 7D, Figs. S9, S10). IFN-γ (essential NK effector), IL-15, 18, BDNF (NK and T cell activation and/or differentiation), RANTES, IP-10, SDF-1α (chemokines associated with NK and T cell recruitment) were among those reduced by anti-CD2 treatment. We also examined changes in genes expression using inflammatory gene expression arrays and found that genes associated with NK and T cell activation and effector function were down regulated following treatment with anti-CD2 (Figs. 7B, 7C, 7E, 7F).

**Figure 7.**
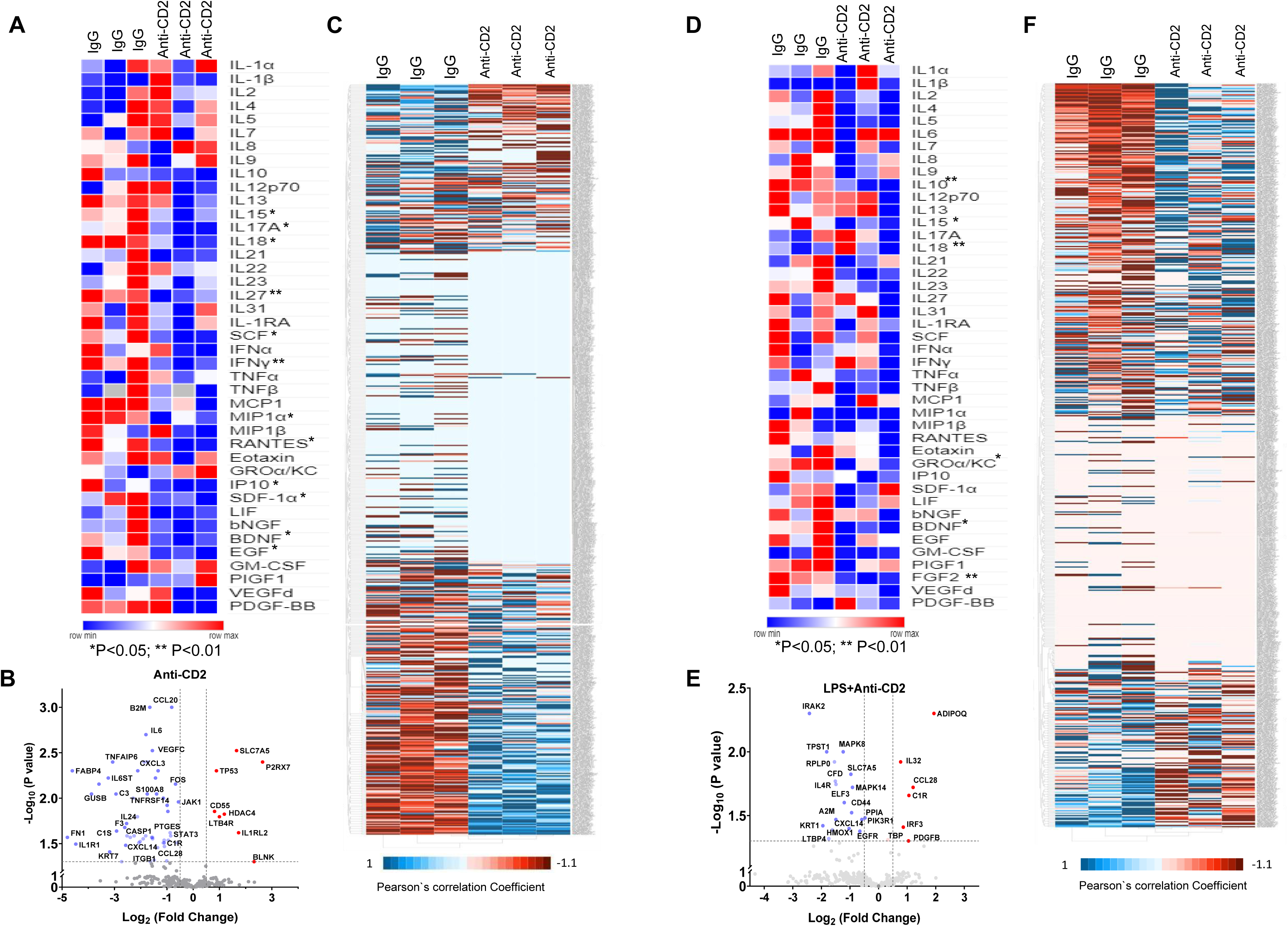
(i): CD2 blockade in organotypic HS skin culture attenuates pro-inflammatory and fibrotic signaling pathways. Skin tissue from HS (n=3) were cultured in transwell plates in the presence of 10µg/mL (**A, C**) anti-CD2 or control IgG antibodies or (**D -F**) LPS (100 nM) with anti-CD2 or control IgG for 72h. (A, D) Student’s t-test of anti-IgG vs anti-CD2 (*P≤0.05, ** P≤0.01). Heatmaps show CD2 blockade decreases the levels of several cytokines/chemokines in culture supernatant relative to control IgG treated tissues. (**B, E**) Heatmaps and (**C, F**) volcano plots depicting changes in gene expression induced by CD2 blockade as determined by TaqMan based human inflammatory OpenArray panel. The data show that CD2 blockade in HS skin organotypic cultures downregulates the expression of several inflammatory and fibrotic genes markers at transcriptional level. The same three skin tissues were used for control-IgG and anti-CD2 treatment.

Anti-CD2 also reversed the pathogenic cytokine expression pattern and gene expression signature in LPS-treated HS skin cultures. LPS is a bacterial component which is known to induce various innate immune responses. Specifically, we found that anti-CD2 treatment led to downmodulation in expression of LPS-induced cytokines/chemokines/growth factors and gene signature associated with TLR2/4 signaling. These include IL-10, IL-15, IL-18, FGF-2, IRAK2, MAPK, ILFR and others (Figs. 7D-F).

We wanted to determine what were the main signaling pathways in HS that are down modulated by CD2 blockade. To address this, we performed ingenuity pathway analyses (IPA) using qRT-PCR HS vs normal skin array data (Fig.1D) and anti-CD2 vs IgG array data (Fig. 7C). The IPA analysis showed several signaling that were upregulated in HS were down regulated by anti-CD2 treatment (Fig. S11A, S11B). The four top ranking upstream regulatory drivers in HS were lipopolysaccharide (P≤8.27E-81, z-score= 6.550), *TNF* (P≤6.13E-70, z-score= 4.994), *IFNG* (P≤3.96E-64, z-score=5.459), and *TGFB1* (P≤1.12E-60, z-score=3.963) (Fig. S12A). Notably, these were also the top four ranking regulatory drivers down modulated by anti-CD2 treatment, *TNF* (P≤1.70E-32, z-score= −3.721), lipopolysaccharide (P≤1.61E-28; z-score= −4.288), *IFNG* (P≤1.72E-27; z-score= −2.993) and *TGFB1* (P≤3.52E-26, z-score= −2.236) (Fig. S12B).

The CD2 interaction with CD58 (LFA-3) leads to bidirectional signals; CD2-signal dependent activation in NK cells and T cells and augmented expression of IFN-γ and CD58-signal co-stimulation in keratinocytes and induced expression of IL-15 and IL-18 (*75*). Thus, a decrease in IL-15 and IL-18 by anti-CD2 treatment is expected. Conceptually, these data reveal a central function for CD2 signaling/co-stimulation in modulating the immunopathogenesis of HS.

## Discussion

Earlier studies report multiple immune cell populations (B lineage cells, plasma cells, neutrophils and proinflammatory macrophages) and molecular mediators (cytokines, complement proteins) are associated with HS pathogenesis. However, the therapeutic interventions based on those discoveries have been minimal (*1*).

In this study, we identified that NKT and NK cell populations intricately involved in HS pathogenesis. Initial analysis of six HS patients by scRNAseq identified cells with a gene signature profile representing NK cell populations (CD3^+^ NKT and CD3^-^ classical NK) are predominant in lesional skin. Other prominent lymphoid cell populations in HS included CD4^+^ T cells and B lineage cells, as previously described (*9, 13, 16, 22*). In HS B cells mostly organized with tertiary follicle or germinal center like structures as recently reported (*76*). Although CD8^+^ T cells have been implicated in HS (*16*), T cells expressing both CD8α and CD8β were underrepresented in our analysis. We further confirmed these data directly by IHC and immunofluorescence staining for CD8^+^ T cells in an independent cohort of HS skin tissues.

CD2 was a highly expressed gene in the scRNAseq dataset of HS patients, which was validated in a cohort of nine HS patients. In fact, it was among the few genes with the greatest increase in expression in HS relative to healthy controls. Remarkably, IHC analysis showed cells expressing high levels CD2 were distributed in all areas of lesional skin tissue and was the most predominant lymphocyte population in HS. In fact, CD2 expressing cells were in far greater numbers than those expressing CD4, indicating they were not CD4^+^ T cells. Many of the CD2^+^ cells expressed the NK marker CD56 that co-expressed CD56, hence NKT cells (*54*). Overall, there were more CD56^+^ cells than CD4^+^ cells. On classical NK cells, CD56 (CD3^-^CD56^dim^) is downmodulated during maturation into a cytolytic population and therefore frequently below the sensitivity of IHC staining (*54*). Nevertheless, we do observe significant numbers of CD3^-^CD56^+^ classical NK cells by IHC.

CD2 is an essential co-stimulatory molecule on NKT cells and NK cells as well as an adhesion molecule through its interaction with LFA3 (CD58) (*43, 44, 47, 56*). The co-engagement of CD2 with CD58 leads to formation of an NK cell immunological synapse, recruitment of Fc-γ receptor, CD16, to the synapse, and generation of signals into the NK cell leading to enhanced NKT or NK cell cytolytic activity and secretion of IFN-γ, the major effector cytokine of cytolytic NK cell populations (*44, 57, 77*). Consistent with this paradigm, we observed that IFN-γ levels were greatly elevated in independent cohorts of HS, as assessed by qRT-PCR and multiplex cytokine analysis. Consistently, elevated expression of IFN-γ was previously reported in other studies (*16, 56*).

The differential spatial distribution of NKT and NK cells also supports the concept that these cells populations have distinct roles in HS pathogenesis. Using CD56 as a defining marker for NK cells, and co-expression of CD3 for NKT cells, we found that CD56^dim^ NK cells were localized in the hypodermal regions and in areas around the periphery of tunnels/sinus tracts. These cells had a highly cytolytic phenotype as they expressed high levels of perforin, granzyme A and granzyme B (*54*). In contrast, CD56^bright^ cells also expressed CD3, a characteristic of NKT, which expressed granzyme A but low levels of granzyme B and perforin. Importantly, they were spatially separated from cytolytic NK cells and physically interacted with fibroblasts in areas of progressive fibrosis as demonstrated by high expression of α-SMA and vimentin. The expression of CD2 was a common feature of NKT, NK cells and CD4^+^ T cells and they interacted with cells expressing CD58 (LFA3), the counterreceptor for CD2 (*43, 44, 56*). Based on morphology and spatial localization, the majority of the CD58 expressing cells in HS were keratinocytes. Since CD58 is expressed widely, including on macrophages, neutrophils, fibroblasts and mesenchymal stem cells, it is possible that their interaction with subsets of NK cell populations contribute to the different aspects of HS pathogenesis. We noted an inverse correlation between CD2 and CD56 expression levels. CD56^dim^ cells expressed higher levels of CD2 and the NK cell maturation marker, CD11b (Mac1), than CD56^bright^ NKT cells and they also contained higher levels of granzyme B. The two cell populations were spatially separated in HS skin. Deeper UMAP sub-cluster analysis of NKT and NK cell population identified six distinct populations based on functional gene expression. Overall, our observations reveal that the NK cell populations in HS are heterogenous and differ in their contribution to the pathogenesis of disease.

HS skin contained elevated levels of CXCL10 (IP10), CXCL12 (SDF-1), CCL1 (MCP1), CCL2 (MIP1-α) and CCL5 (RANTES) chemokines that have been shown to be major factors required to recruit NK cells to tissues. In the skin they have been observed in inflammatory diseases such as psoriasis (*78*). These chemokines are produced largely by keratinocytes when activated by the inflammasome pathway. TLR signaling and downstream activation of the NLRP3 inflammasome is a feature of HS pathogenesis (*18, 20*). In addition, we also observed increased levels of IL-12, IL-15 and IL-18, the cytokines that individually or in combination activate NK cells and enhance their proliferation and functional activity (*52, 72*). In the skin, these cytokines are primarily produced by keratinocytes following activation of TLR-inflammasome pathway. In HS, we found elevated levels of IL-8 (CXCL8) which is the major chemokine that recruits neutrophils to sites of inflammation (*79*). Although, we do not know which cell type/s produce IL-8 in HS, this chemokine is produced by CD4+ T cells, NK cells, keratinocytes, and macrophages in inflamed tissue (*79*).

As discussed above, the transmembrane glycoprotein CD2 is an adhesion and co-stimulatory molecule that is expressed on T cells, NKT cells and NK cells and interacts with CD58 (LFA3) expressed on antigen presenting cells and other cell populations including keratinocytes (*57*). The cognate interaction of CD2 with CD58 contributes to the generation of an immunological synapse and provides essential costimulatory signals for optimal activation, differentiation/maturation and proliferation of T cells and NK cells (*44, 56, 75*). Blockade of CD2-LFA3 engagement is immunoregulatory and shown to be important for suppressing immune responses particularly in the transplantation setting (*43, 80*). Siplizumab is an anti-CD2 mAb that has received FDA approval to prevent transplant rejection (*80*). Since we found elevated CD2 on NK cell populations and CD4^+^ T cells in HS, we tested if its blockade has therapeutic potential for the disease. It should be appreciated that there is no relevant animal model for HS, therefore we tested blockade of CD2 using anti-CD2 mAb in our newly developed organotypic culture system (*74*). Treatment of HS tissue with anti-CD2 mAb from three different individuals with anti-CD2 mAb led to dramatic downmodulation of many genes associated with immune and non-immune cell activation, inflammation, and proliferation. Expression of genes, such as *CD55, TP53* (p53), important for enhancement of tolerogenic activity, suppression of NK cell function and/or attenuating inflammation were increased in anti-CD2 mAb treated HS tissues (*81*). Anti-CD2 mAb treated tissues also showed a significant decrease of several inflammatory cytokines and chemokines compared to control IgG treated cultures. The outcome of anti-CD2 mAb treatment was similar in HS tissues that were stimulated with LPS. Thus, CD2 serves as an important therapeutic target for the diminution of painful inflammatory response in HS patients.

Our miRNA profiling assay revealed that eleven of the 36 miRs differentially expressed in HS vs controls, have previously been shown to be important for NKT and NK cell differentiation and/or function (*61, 62*). Among them was miR-155 that enhances expression of IFN-γ in NK cells and is induced by IL-12 and IL-18 (*82, 83*). In HS miR-155 was highly elevated (12.5 fold) compared to controls and so was the expression of IL-12 and IL-18. Some miRNAs have opposing biological effects in NKT and NK cells during immature vs mature stages of differentiation. For example, miR-150 which is necessary for development and differentiation of NK cells into mature NK cells represses generation of NKT cells (*61*). At sites of inflammation, miR-150 regulates expression of perforin (*61*). Previous studies have shown that micro RNAs, miR-27a-5p and miR-181a, attenuate NKT cells or NK cells differentiation (*84*). In HS, we found that these miRs were expressed at lower levels than in controls. The composite data supports a regulatory network favoring NKT and NK cell expansion and activation in HS pathogenesis.

In summary, we report a novel role for innate immune lymphocytes, specifically NKT cells and NK cells, in HS pathogenesis that may be the “missing link” as to why most treatments have had limited benefit. Our studies reveal a striking heterogeneity in NKT and NK cell populations in HS skin, with each subpopulation contributing to mostly non-overlapping aspects of disease pathogenesis as depicted in **Fig. 8**. A common feature of all NKT and NK cell populations was the elevated expression of CD2, an adhesion and activating receptor, that is integrally involved in the cellular network of HS pathogenesis through its interaction with CD58 on keratinocytes and fibroblasts. Disruption of this interaction using anti-CD2 blocking antibody attenuated HS pathogenesis associated cytokine/chemokine protein and gene expression. We recognize that the HS tissues studied here were mostly from patients who were refractory to all previous treatments. Therefore, the striking contribution of NK cell populations may reflect such a patient population. Nevertheless, CD2 blockade may be beneficial in all stages of HS.

**Figure 8:**
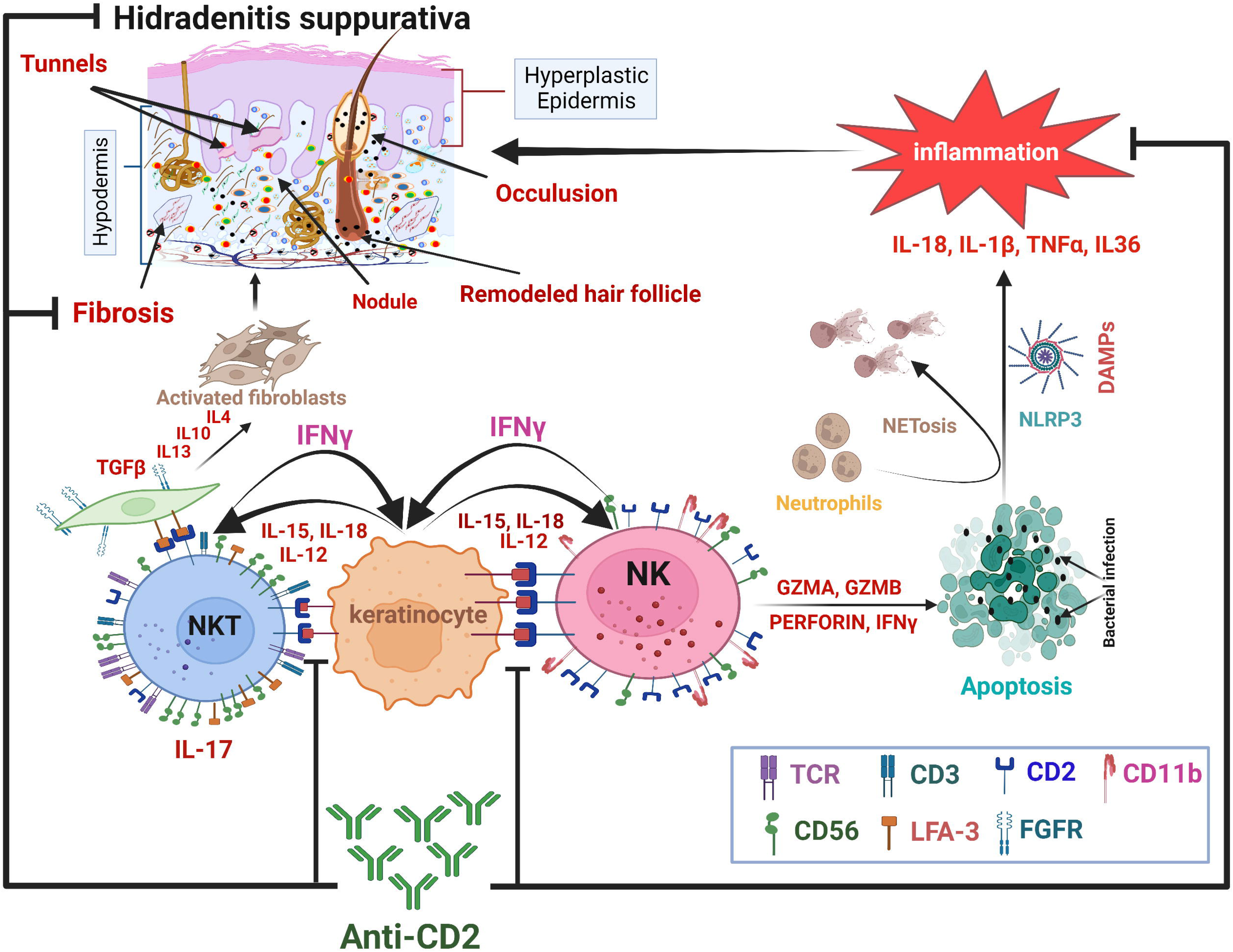
CD2 blockade attenuates NKT and NK cell activities in HS pathogenesis. Diagrammatic representation of NKT and NK cell populations in HS pathogenesis and role of CD2. Subpopulations of NKT cells and NK cells drive tissue destruction and remodeling in HS. CD56^bright^ NKT cell subpopulations expressing IFN-γ, TGF-β, IL-13, IL-10 and/or IL-4 interact with keratinocytes and/or fibroblasts. CD11b^+/-^ CD56^dim^ NK cells producing IFN-γ interact with keratinocytes. Cognate interaction of CD2 on NKT and NK cell populations with CD58 on keratinocytes, fibroblasts, and innate immune cells (not shown) is critical for NKT and NK cell dependent disease processes.

## Material and Methods

### Human Subject

The Institutional Review Board of the University of Alabama at Birmingham approved the protocol (IRB-300005214) for obtaining surgically discarded skin tissues from healthy and HS subjects. Surgical excised skin from 24 patients with HS (Hurley stage II or III; 22 Black (18 males, 3 males, 1 unknown) Caucasian male (1), Unknown ethnicity (1) (Table S7). All were >18 years of age. 19 normal skin from breast or abdominoplasty reduction surgery (13 Black females, 2 Caucasian female, 3 Black males and 1 Hispanic male; age> 18 years), was used in the study. Fresh tissues were processed for single cell RNA Seq and *ex vivo* skin explant cultures.

### Single Cell Preparation

Skin (3×3 cm) was placed in 5 ml of medium 154 supplemented with human keratinocyte growth supplement and cut into small pieces. Freshly prepared 100µl Liberase TL (2mg/mL stock, Roche Diagnostics) + 700μl of Tyrode’s solution was added to the tissues and incubated for 15 min at 37°C and tissue was dispersed through a 70μm cell strainer (Falcon).

### Single cell RNA sequencing

10X Genomics protocol was used for generation of libraries for scRNA-seq. Normal skin scRNAseq data were from two in-house and two public data sets (GSM6840146 and GSM6840150) (*85*). 10x Genomics Cell Ranger (v.5.0.1) tool with human reference was sued for data processing, including quality control, read alignment, and gene quantification. Seurat (v.4.0) package was used for tertiary analysis of scRNAseq data (*40, 86*). Cells with the parameter (percent.mt < 25, nCount_RNA < 60000 & nFeature_RNA < 6000) and doublets/Multiples cells (Chord tool (*87*)) were identified and removed using a highly optimized machine learning tool. Log normalization was used to normalize the scRNA-seq data counts. To reduce the impact of batch-effect we used harmony (*41*) package for integration of data. For cell clustering 20 dimensions and 0.9 cluster resolution was used. We identified 23 clusters by the UMAP dimensional reduction algorithm. The Azimuth based markers along with previously reported strategy for HS scRNAseq cell type identification was used annotation of cell populations (*16*). The markers genes used for annotation are included in the dot plot. For the sub-clustering analysis of immune cells 30 dimensions and 0.7 resolution was selected. Cell-type specific gene expression analysis was performed by FindAllMarkers function with parameters min.pct = 0.25, logfc.threshold = 0.25, and slot = “data”.

### Bulk transcriptomics data acquisition and analysis

To study the bulk transcriptome expression profiles altered by HS, we extracted four publicly available datasets (GSE151243, GSE154773, GSE79150, and GSE128637) representing both microarray and RNA-sequencing (RNA-Seq) technologies form NCBI GEO database (*88*). The Raw values were log_2_ normalized for further analyses hereafter. The differentially expressed genes (DEGs) analysis for each dataset was performed by DESeq2 (*89*) for RNA-Seq and edgeR (*90*) for microarray experiments with default parameters (FDR<0.05, log_2_FC =|1|).

### Pathway enrichment analysis

The canonical pathway analysis on DEGs was performed using Ingenuity pathway analysis (IPA)(*91*) with significant fold change and test statistics values. Additional function enrichment analysis of bulk transcriptome was performed by enrichR (*92*) for KEGG pathway analysis platforms with standard significance (*q*-value <0.05).

### Visualizations

Single cell RNA seq analysis plots (clusters, heatmaps, dot plot, violin plots) were generated through R (v4.0.3) statistical software. All experimental verifications’ violine plots, volcano plots and box plots were drawn using GraphPad Prism.

### Statistical analysis

To identify the marker genes in single cell RNA-seq clusters, a non-parametric Wilcox test was used with default parameters. To identify the DEGs in bulk transcriptome datasets a standard cutoff (*fdr*≤0.05; *log2fc* ≥|1|) was used. The activated and inhibited canonical pathway analysis was determined by parameters (*z-score* ≥|1|; *BH p-value* ≤*0.05*). The significance of experimental verified DEGs was (students T. test (*p-value*) ≤0.05 and *log2fc* ≥|1|). The functional enrichment of genes/proteins was performed with significance (-log10(*q-value* > 1.3). To identify the DEMs in miRNAome datasets a standard cutoff (*p-value*≤0.05; *log2fc* ≥|1|) was used.

### Quantitative qPCR Open Arrays

To validate global differential genes expression from the publically available RNASeq datasets of normal and HS skin samples, we employed quantitative TaqMan based real-time qPCR for predesigned gene panels of various regulatory pathways. We used human Inflammation open array panel (Cat # 4475389), human signal transduction panel (Cat # 4475392), human kinome panel (Cat # 4475388) and human stem cell openarray panel (Cat # 4475390, ThermoFisher, USA). Briefly, total RNA from HS and normal skin tissue was isolated using Trizol reagent (Cat # 15596018, ambion). A total of 2µg of RNA was reverse transcribed into cDNA using SuperScript® VILO™ cDNA Synthesis Kit (Cat #11754250, Life Technologies). Pre-amplification of cDNA was performed as per the manufacturer (Thermofisher). The pre-amplification products were processed as per the manufacturer’s instructions. OpenArray chips in total contained primers for 2429 targets were read on 12_K Flex RT-PCR machine (Thermo Fisher Scientific, Life Technologies Corporation, Grand Island, New York). Data analysis was performed through the online available Expression suit v 1.3 software (Thermo Fisher Scientific, Life Technologies Corporation, Grand Island, New York) using global gene normalization method.

For microRNAs profiling, we used quantitative TaqMan based OpenArray Human MicroRNA Panel (Cat# 4470187; Thermo Fisher Scientific, Life Technologies Corporation, Grand Island, New York). This panel contains 754 well-characterized human miRNA sequences from the Sanger miRBase v14. All assays were performed as manufacture’s instructions. Data analysis was performed through the online available Expression suit v 1.3 software (Thermo Fisher Scientific, Life Technologies Corporation, Grand Island, New York) using global gene normalization method. Bioinformatics analysis of all panels was carried out using Ingenuity Pathway Analysis (Qiagen, Version 84978992).

### Sample Processing and *Ex Vivo* Culture

Clinical specimens were divided into ∼5 mm x 5 mm tissue pieces. Tissue pieces were then placed on to a 0.4 µm transwell filter (Merck Millipore) with a thin layer of extracellular matrix (ECM, 90% collagen type 1 (Advanced Biomatrix, USA) + 10% growth factor reduced Matrigel (Corning, USA) for stability as described previously (*74*). KBM Gold Basal Medium (00192151; Lonza) containing KGM Gold supplements (00192152, Lonza) was added to the transwell filters and to the bottom of the well to generate air-liquid (the media volume did not cover the top tissue layer) cultures. Cultures were treated with anti-human CD2 mAb (10µg/mL; Cat# 300240; Clone: RPA-2.10; Biolegend) or IgG (10 µg/mL, Cat# 403502, Clone: QA16A12, Biologened) 72h. The supernatant culture media was collected daily and stored at −80C for profiling of cytokines/chemokines/Growth factors.

### Multiplex cytokine/chemokine assay

We used the cytokine/chemokine/growth factor 45-Plex Human ProcartaPlex™ Panel 1 (Cat# EPX450-12171-901, ThermoFisher, USA) was used. Briefly, the tissues were homogenized using cold RIPA buffer (Santa Cruz biotechnologies) containing 2 mM sodium orthovanadate, 1 mM PMSF, and protein cocktail inhibitor (1X; Santa Cruz). The measurement were performed suing Luminex 200 instrument (Luminex Corporation, USA) as described (*93*).

### Histological Analysis and Confocal Immunofluorescence (IF) staining

#### Hematoxylin and Eosin (H & E) staining

Briefly, skin tissues were fixed in 10% formalin, embedded in paraffin and 5_μm sectioned were prepared. The skin sections were deparaffinized in xylene, rehydrated and stained with H& E. Images were captured using BZ-X710 bright field microscope and integrated for Z-stacking and stitching with BZ-X analyzer (Keyence Corporation, Japan).

#### IHC Staining

For IHC analysis, serial tissue sections (5-μm thickness) were cut from FFPE tissue blocks of human HS (n=5-7) and normal skin (n=3) biopsies using a microtome and mounted on poly-L-lysine–coated silicon substrate for immune cells analysis. Sections were baked at 65 °C for 15 min, deparaffinized in xylene (3X 5min) and rehydrated via a graded ethanol series (100% ethanol for 5 min twice, 95% ethanol for 5 min, 75% ethanol for 5 min, 50% ethanol for 5 min followed by ddH2O for 5 min). The sections were then immersed in Antigen unmasking solution (Vector, H3300) and placed in a microwave at power 10 for 2 min followed by 8 min at Power 3. The sections were subsequently rinsed twice with dH_2_O and once with wash buffer (TBS, 0.4% Tween 20, pH 7.2). Residual buffer was removed by gently touching the surface with a lint-free tissue and hydrophobic barrier was created around the specimen using pep pen and Bloxall blocking solution (SP-6000) was used to block endogenous peroxidase activity. Sections were then incubated with blocking buffer for 30 min (TBS, 0.4% Tween, 3% BSA, 10% Normal Goat Serum, pH 7.2). The sections were incubated in various antibodies against CD2 (1:1000, ab131276, Abcam), CD56 (1:500, ab9272, Abcam), CD3 (1:100, ab5690, Abcam), CD4 (1:500, 133616, Abcam), CD20 (1: 100, ab78237, Abcam) and CD8β (1:250, PA5-80450ThermoFischer Scientific) overnight at 4 °C in a humidified chamber. HRP conjugated goat anti-rabbit/ goat anti mouse (1:200, abcam) were used as secondary antibodies. DAB Substrate Kit (Cat# ab94665, Abcam) was used for amplification and visualization of signal, respectively. Images were captured and stitched at 20x magnification using BZ-X710 Keyence microscope for entire area of tissue. We used Macro Cell Count module of BZ-X analyzer which measure multiple files (upto 400) in a single operation according to the settings recorded in the condition file. Debris/cells having size less than ≤4μM of size were removed from analysis. The total area of all images was also calculated using BZ-X analyzers. Unpaired ‘t’ test with Welch’s correction was used to identify statistical significance (0.05≤P) between experimental groups.

#### Immunofluorescence Confocal Analysis

skin sections were deparaffinized, rehydrated and then incubated in antigen unmasking solution according to the manufacturer’s instructions (Vector laboratories, Burlingame, CA). Sections were blocked in blocking buffer containing 5% normal goat serum in PBST (PBS+0.4% triton X100) for 1h at 37_°C. Sections were then incubated with primary antibodies against various proteins in blocking solution for overnight at 4_°C; antibodies listed in **Supplemental Table 8**. Sequential staining was done for visualization of more than one target in a single specimen. After washing (3X times, 10 min each) with PBST, sections were re-incubated with various fluorescence-coupled secondary antibodies (1:200, Invitrogen). Sections were fixed in DAPI containing Vectashield gold antifade medium (Cat#H-1200, Vector laboratories, Burlingame, CA). Sections were then visualized under FLUOVIEW FV3000 confocal microscopes (Olympus, USA) equipped with FV3000 Galvo scan unit. Z-projected images were post-processed for noise reduction, 3-D image construction and movie preparation using FV3IS-SW version 2.3.2.169 software provided with Olympus Fluoview F3000 confocal microscope. OLYMPUS cellSense Dimension (version 2.3) was used for counting the cells numbers at various locations and samples. Ordinary one-way ANOVA multiple comparisons (Two stage linear step-up procedure of Benjamini, Krieger and Yekutieli) was used to determine statistical significance among the groups.

#### TUNEL assay

Apoptotic cells were detected using In Situ Cell Death Detection Kit, Fluorescein (Cat#11684795-910, Roche, Germany).

## Author contributions

Conceptualization and experimental design by M.P.K., B.M., M.A., M.S.M. and C. R. Majority of experiments were performed by M.P.K., and scRNAseq, bulk RNA-Seq, and systems biology by B.M. Additional authors including R.S., L.J., N.K., and K.G. provided technical input and assisted with experiments. Initial data evaluation by M.P.K., B.M., M.A., M.S.M and C.R. Clinical evaluation of patients and tissue procurement: B.E.E and C.A.E. Writing-original draft: B.M. C.R. and M.P.K. Writing-final draft: C.R., M.P.K, M.A., B.M. and M.S.M., Editing and critical comments by C.A., B.E.E. and J.D.

## Funding

This work is supported by NIH/NIEHS grant R01 ES026219 and NIH/NCI grant 5P01CA210946 to M.A., and R21 AI149267, R21 MDO15319 to C.R.

## Competing interests

The authors declare that they have no competing interests.

## Data and materials availability

The processed data of this study have been deposited in the GEO database under accession code. All data, codes, and materials used in the analysis are available within the Article, Supplementary Materials or from the corresponding author upon reasonable request.

## Notes

### Competing Interest Statement

The authors have declared no competing interest.

