## Supplemental Figures for "NK and NKT cells in the pathogenesis of Hidradenitis suppurativa: Novel therapeutic strategy through targeting of CD2"

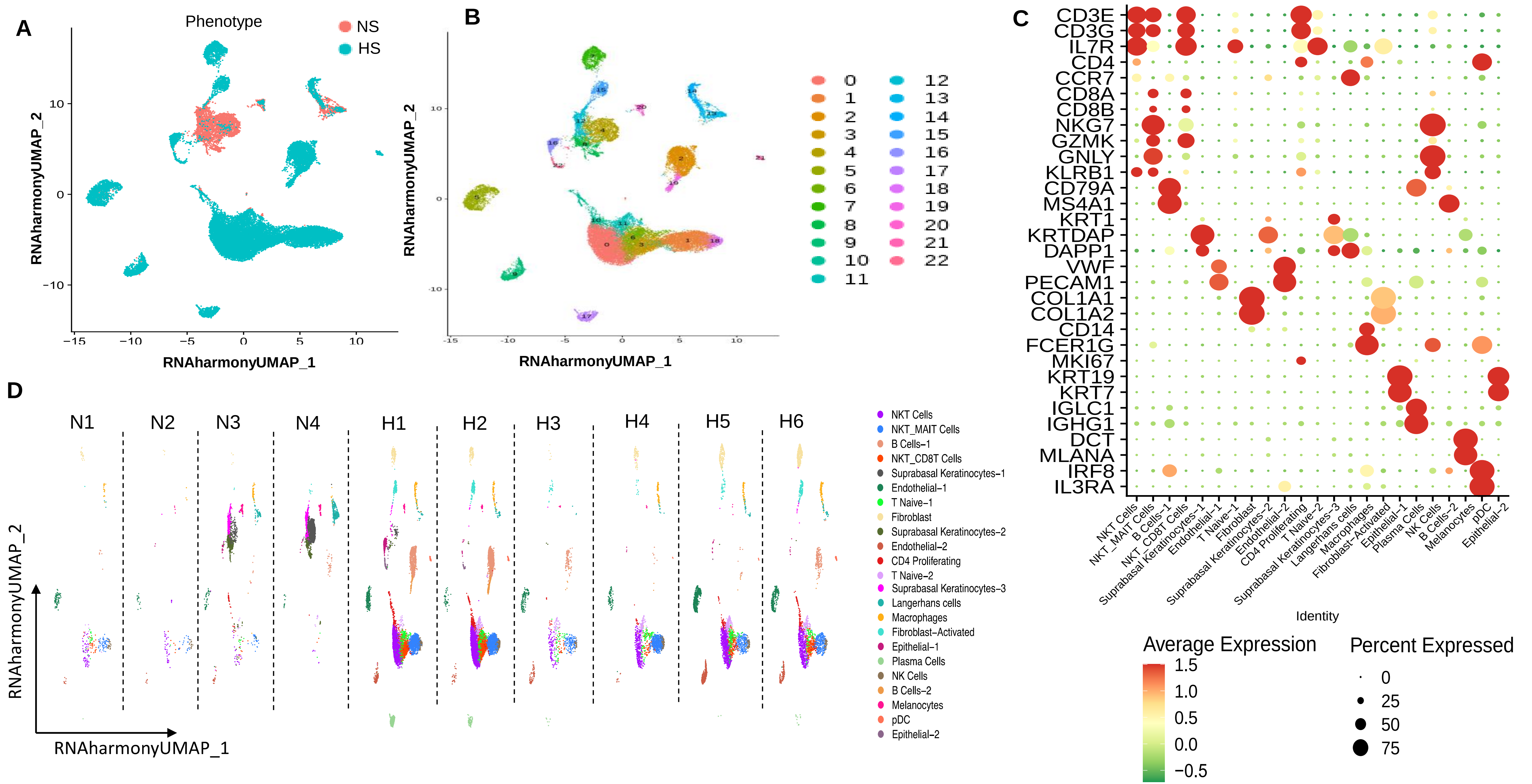


**Figure S1. Annotation of different cell types based on re-clustering and unsupervised annotation**. The normal and HS samples met QC, were used in subsequent annotation of various clusters. **(A)** Graph depicting various cell clusters in NS (Coral color) vs HS (Cyan color) samples. **(B)** Graph showing 23 different clusters. **(C)** Cell biomarkers used for cluster designation. (D) Graphs showing various cell clusters in each individual samples.

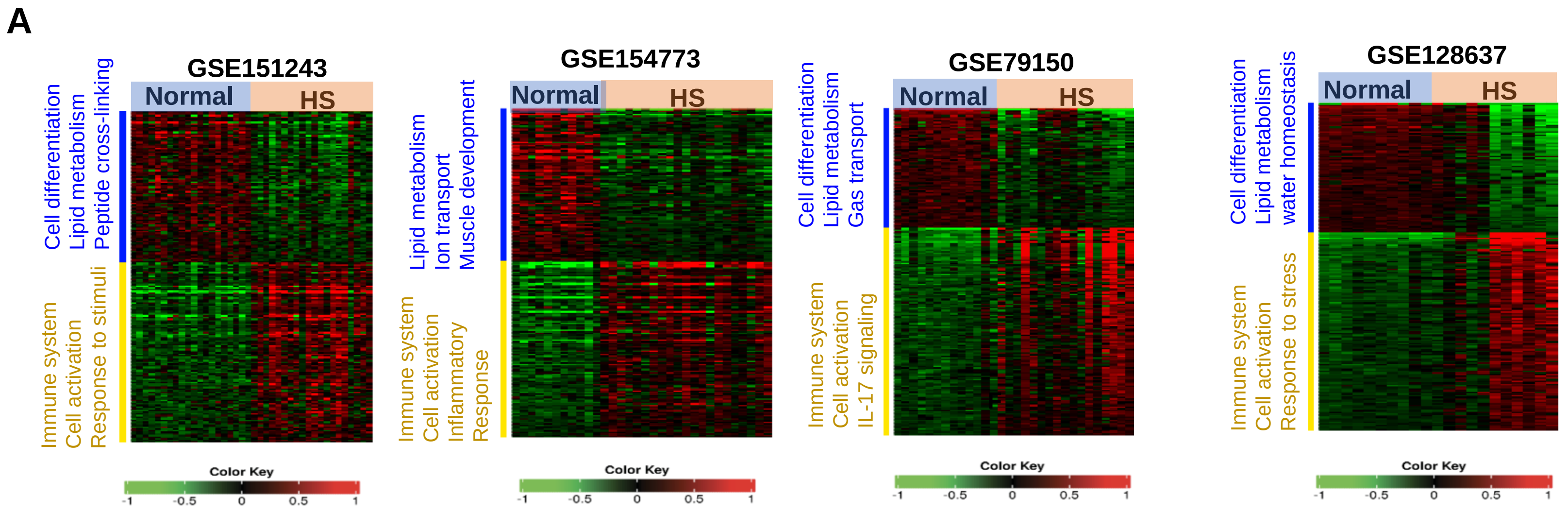


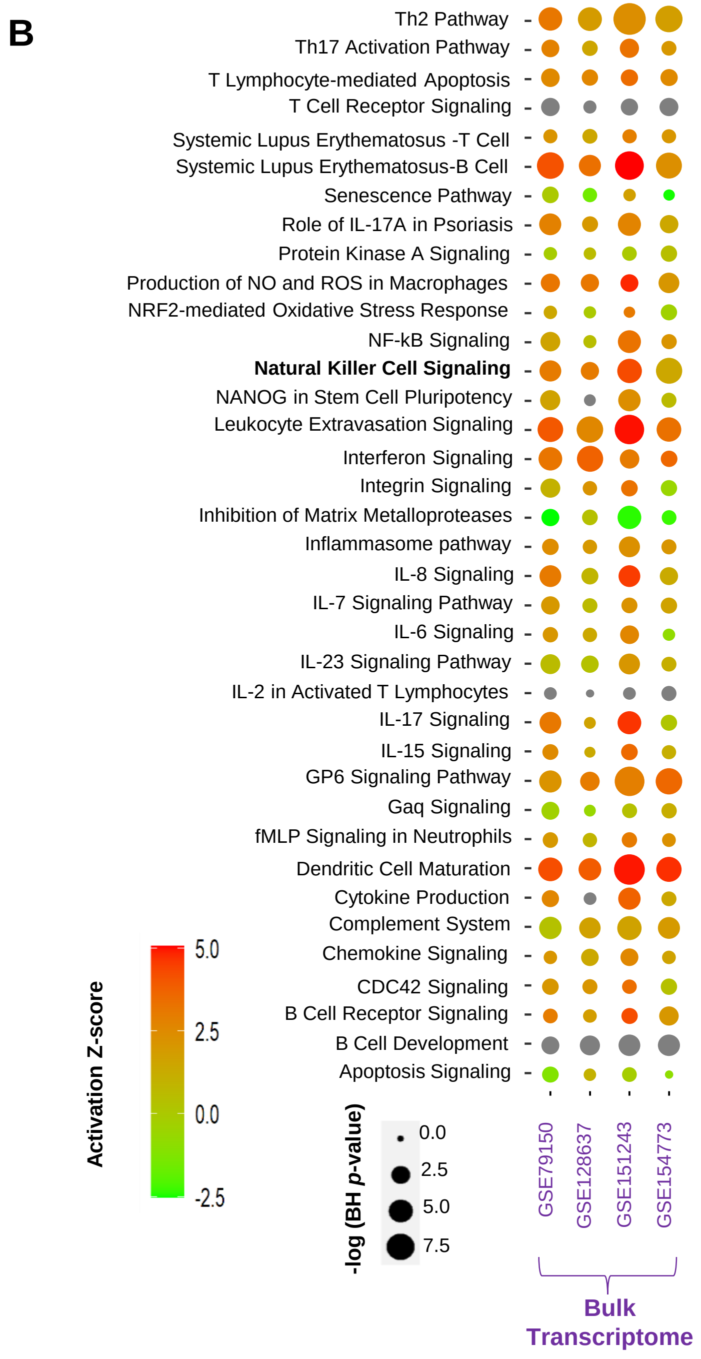


**Figure S2. Bulk transcriptomics analysis .** (**A**) Heat map representing bulk transcriptomic analysis of four public available datasets (GSE151243, GSE154773, GSE79150, and GSE128637) generated from both microarray and RNA-sequencing technologies. Data show comparison between controls (normal/non-lesional) and HS (HS or lesional skin). **(B)** The significant activated or inhibited canonical pathways in each dataset identified using IPA (-log (BH p-value ≤0.05; z-score > |1|, grey dots are enriched pathway.

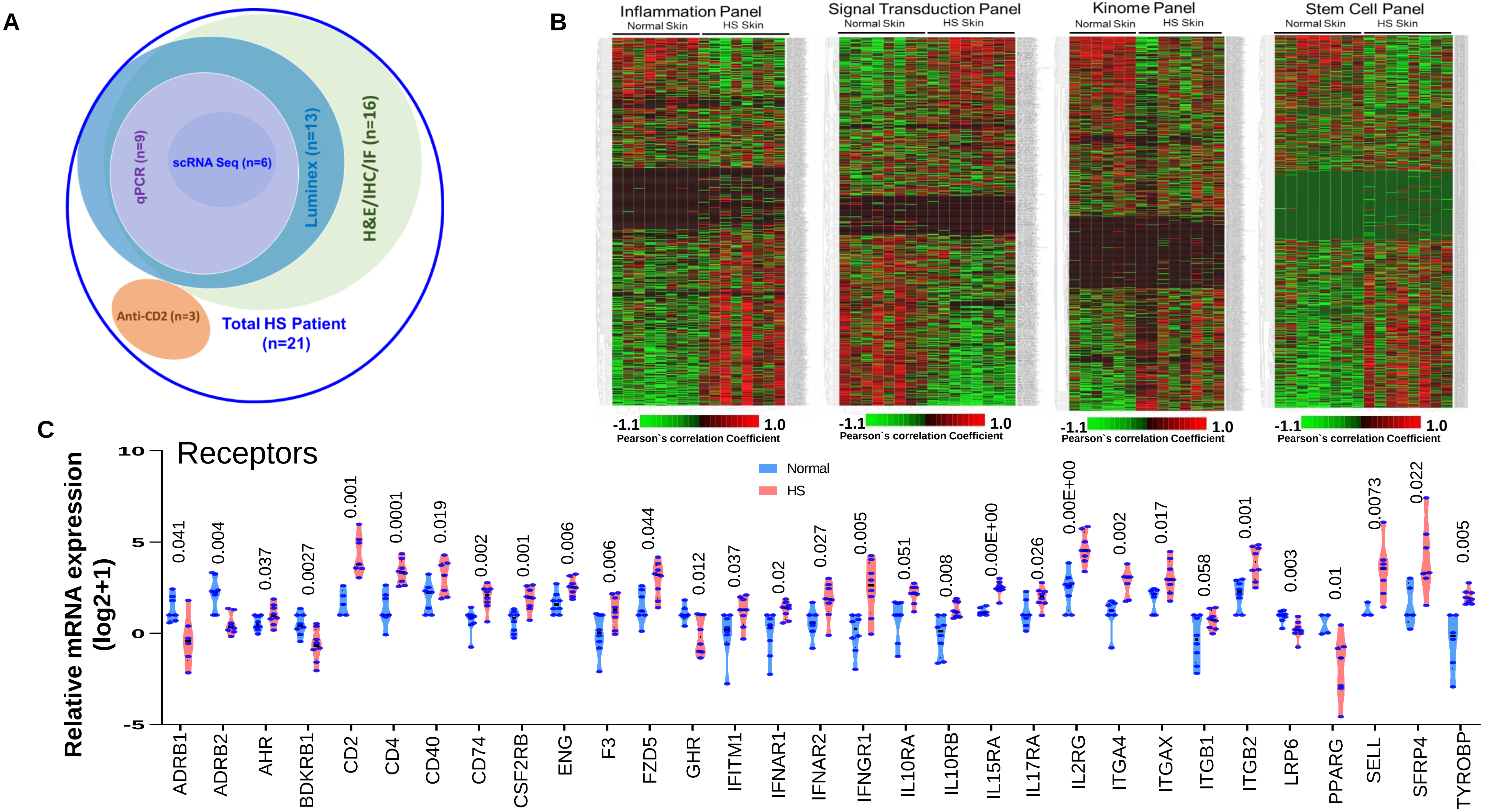


**Supplementary Figure S3. qRT OpenArray PCR analysis: (A)** Venn diagram showing skin samples from HS patients used in various experiments. Overlapping circles represent patient`s sample used in more than one experiment. **(B)** Heatmap of qRT-PCR OpenArray analysis of four different signaling panels containing 2429 unique target genes in skin samples from normal (n=6-8) and HS (n=8) subjects. **(C)** Violin plots showing significantly changed (P≤0.05) in the mRNA expression of various receptors in HS vs NS

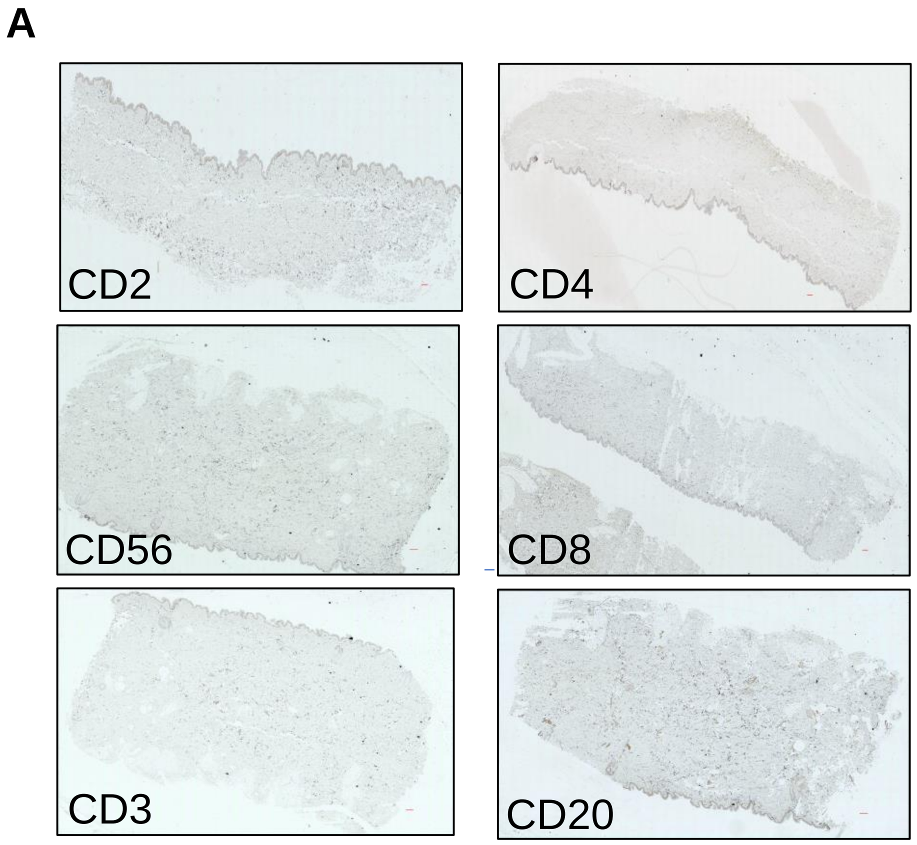


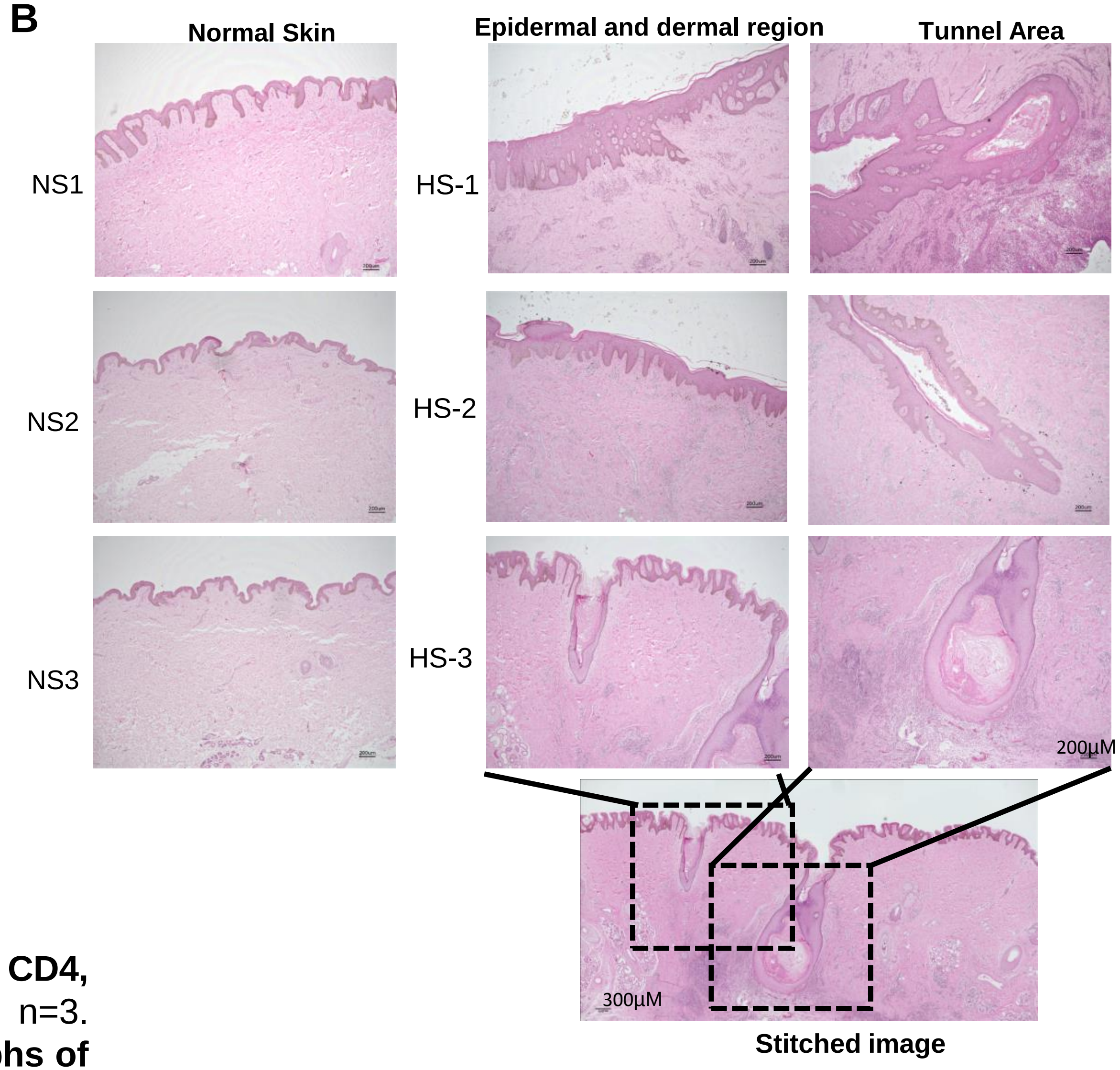


**Figure S4. A. IHC of normal skin stained for CD2. CD56, CD3. CD4, CD8 and CD20 expressing cells.** Figures are representative, n=3. Small red bar in each panel = 300 µM. **B. H&E stained micrographs of normal (n=3) and HS (n=3) skin samples**. The images show morphological anomalies in HS skin - epidermal hyperplasia, leukocytes hyper-infiltration and tunnels. (Magnification= 4x, Scale bars= 200µm and 300µM for stitched image).

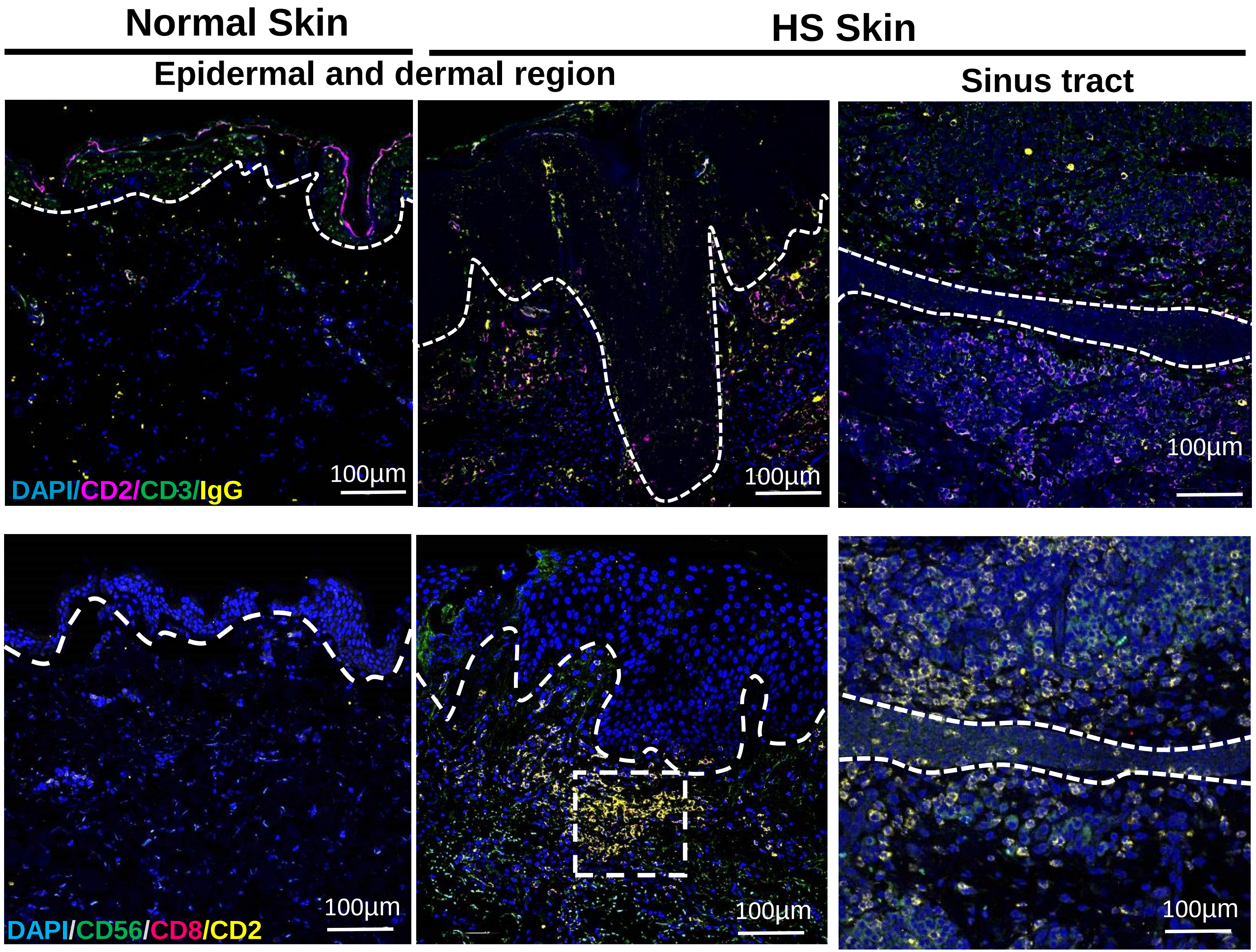


**Figure S5. Plasma cells and CD8β cells in HS.** ***Upper panels*****:** Micrographs showing immunofluorescence staining for IgG (plasma cells). ***Lower panels*****:** Micrographs showin immunofluorescence staining for CD8 β in HS. Resolution 4096X4096, Magnification= x20, Scale bars= 100µm.

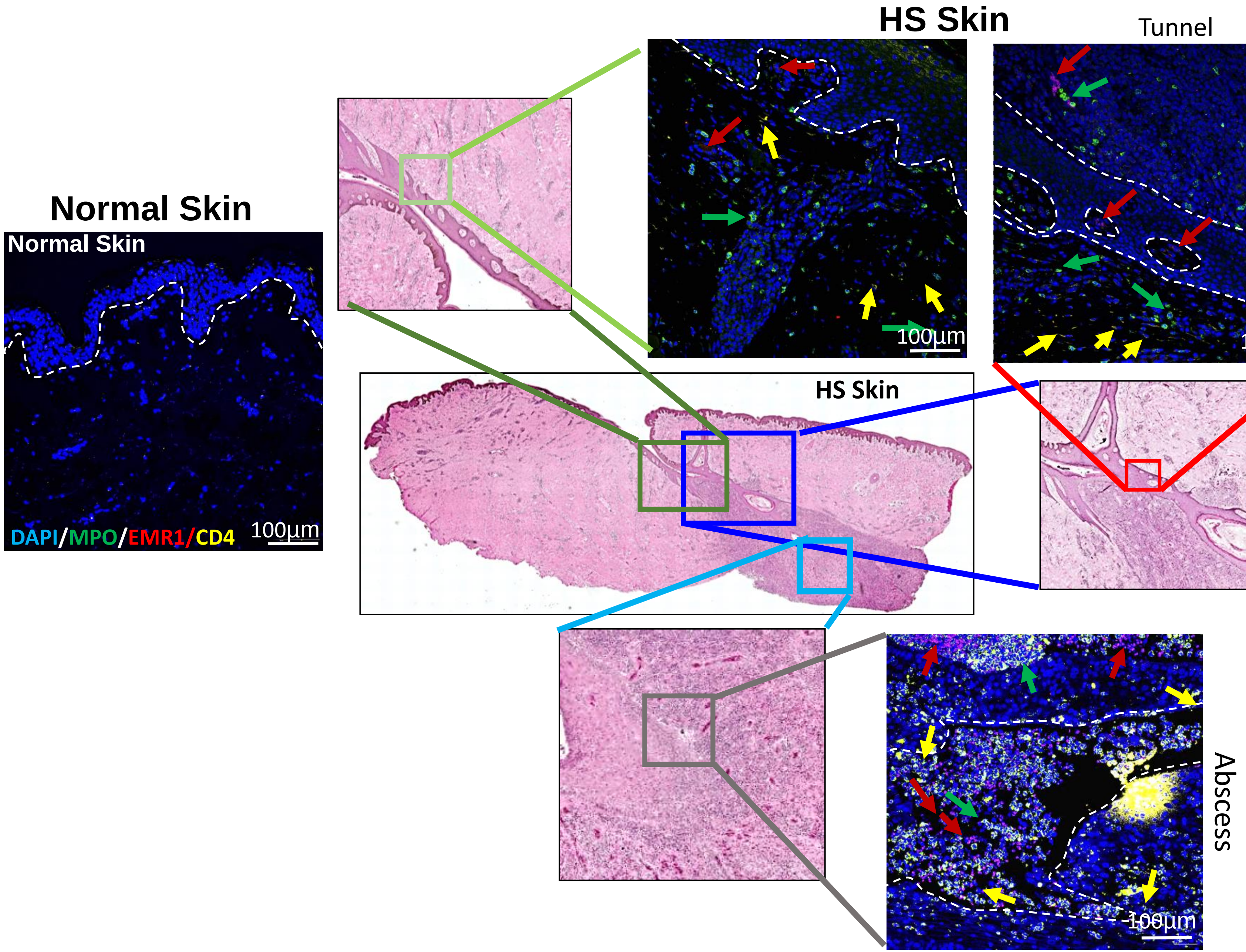


**Figure S6. Neutrophils and macrophages in HS skin.** Micrographs for immunofluorescence staining for MPO (neutrophils), EMR1 (macrophages) and CD4+ T cells in normal and HS (epidermal/hypodermal, sinus tract, abscess). Arrow by color designate cell populations. (Resolution 4096X4096, Magnification= x20, Scale bars= 100μm)

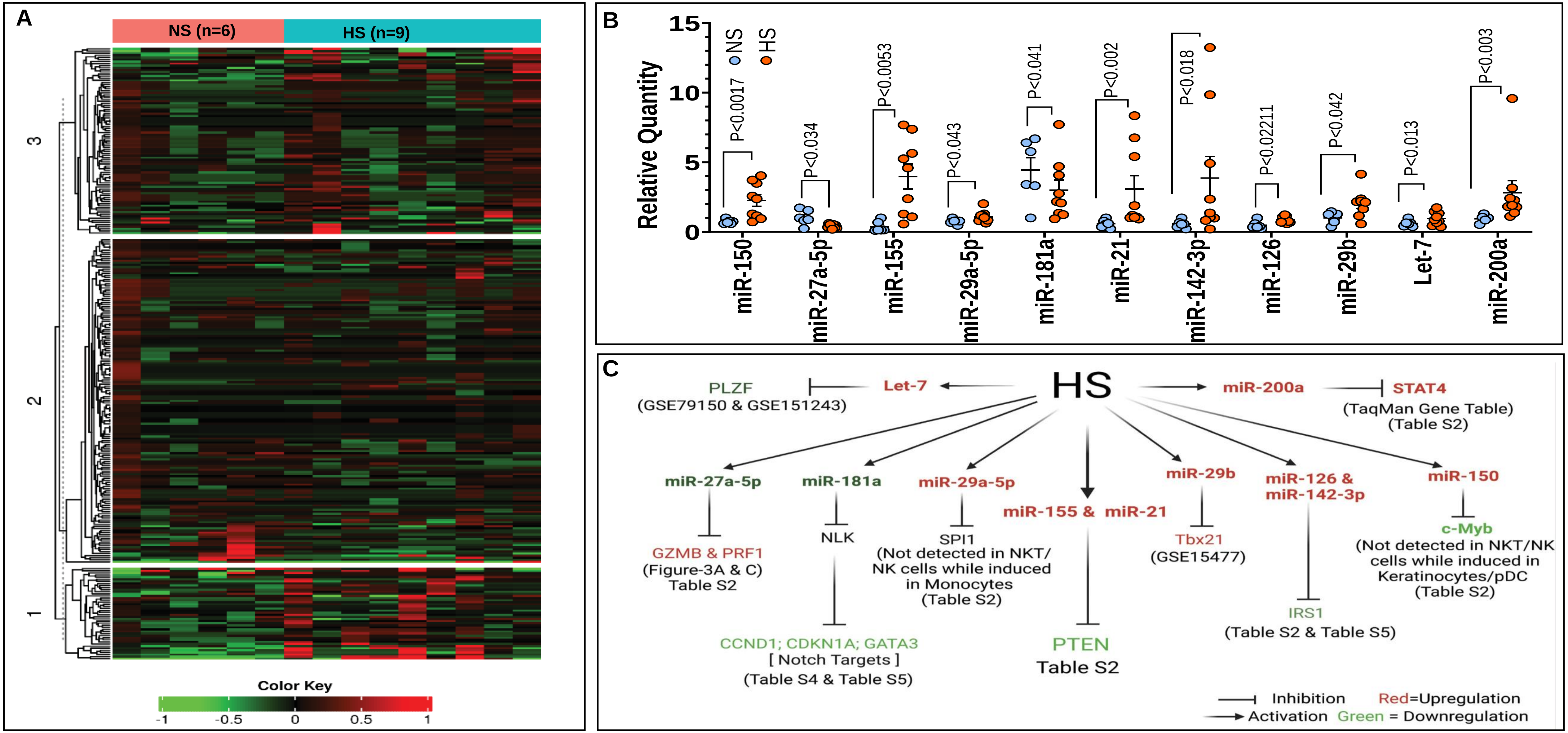


**Figure S7. miRNA profiling in normal and HS samples. (A)** Heatmap of expressed miRNAs in normal and HS skin samples. miRNA profile in skin from NS (n=6) and HS (n=9) were determined using OpenArray panel containing 754 well characterized miRs. **(B)** Dot plot showing relative quantification of eleven miRs with known role in modulating NKT/NK differentiation/maturation/function. Each dot represents tissue from an independent NS or HS individual. **(C)** Diagram depicting downstream gene targets of miRNAs modulated in HS with relevant function in NKT and/or NK cells.

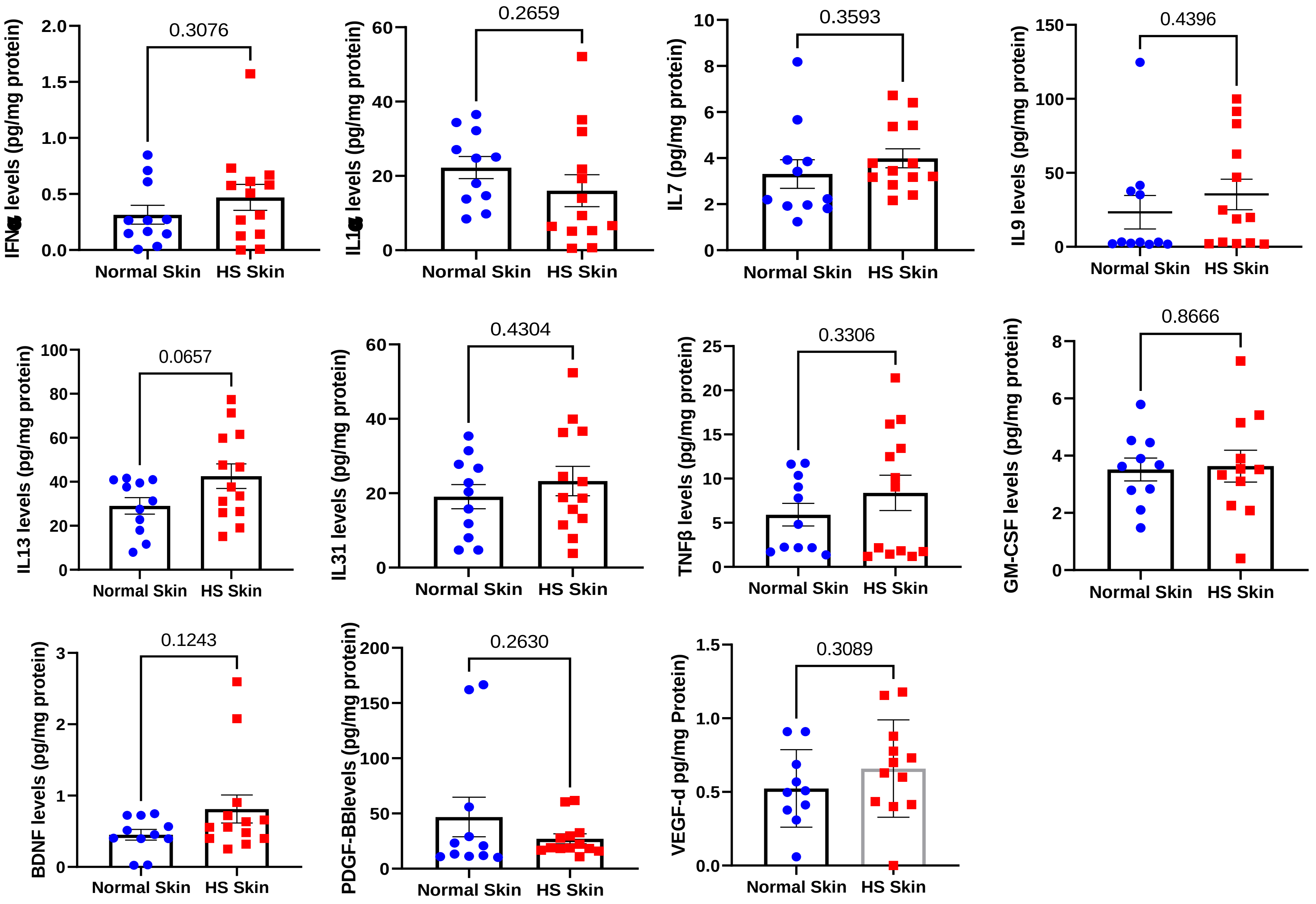


**Figure S8. Cytokine, chemokines and growth factors in NS and HS.** The data are analytes in which levels in HS were similar to NS. Each dot represents an individual (NS=11, HS=13)

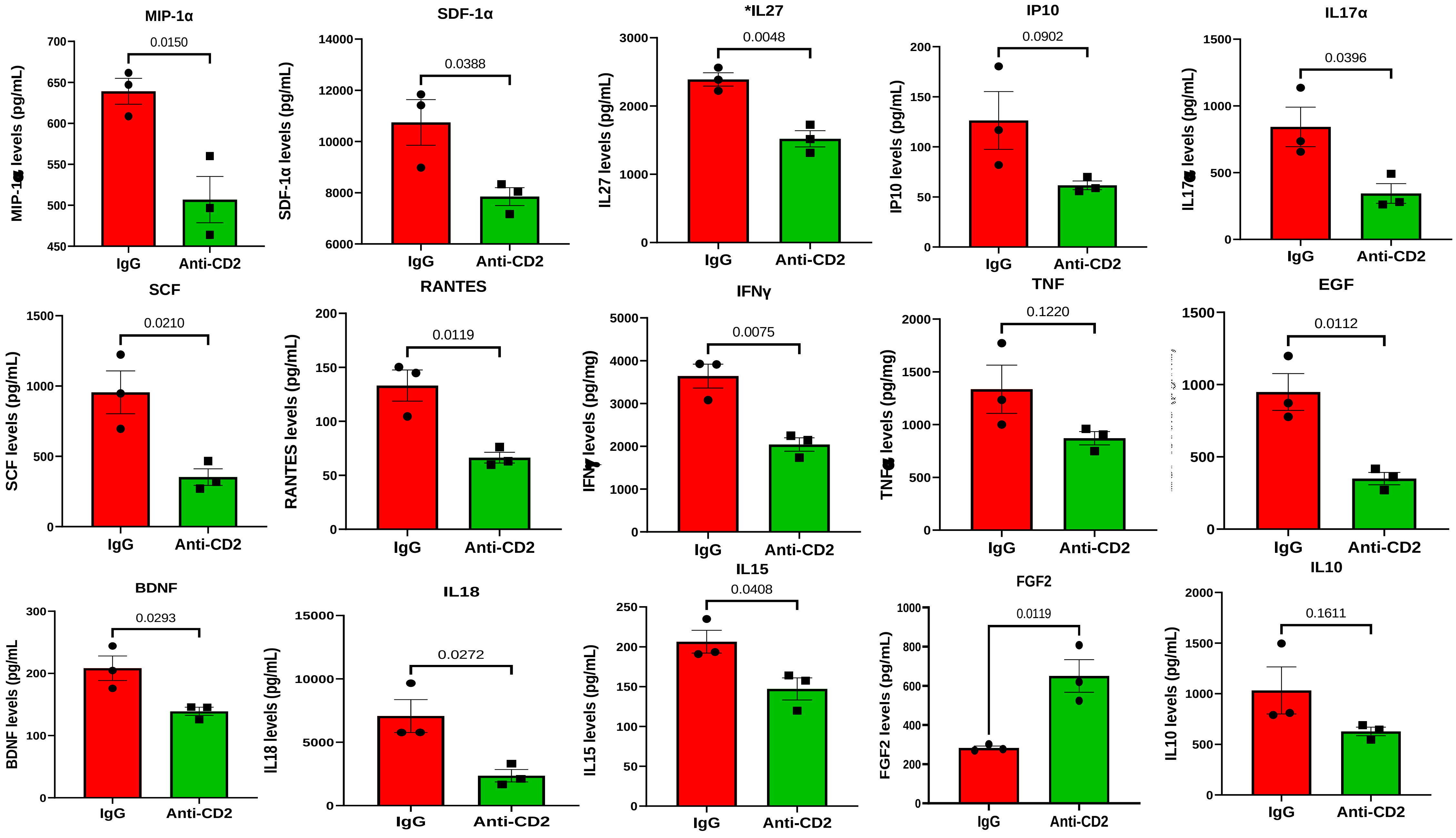


**Figure S9. Levels of cytokines, chemokines and growth factors in anti-CD2 or IgG treated organotypic cultures of HS skin.**

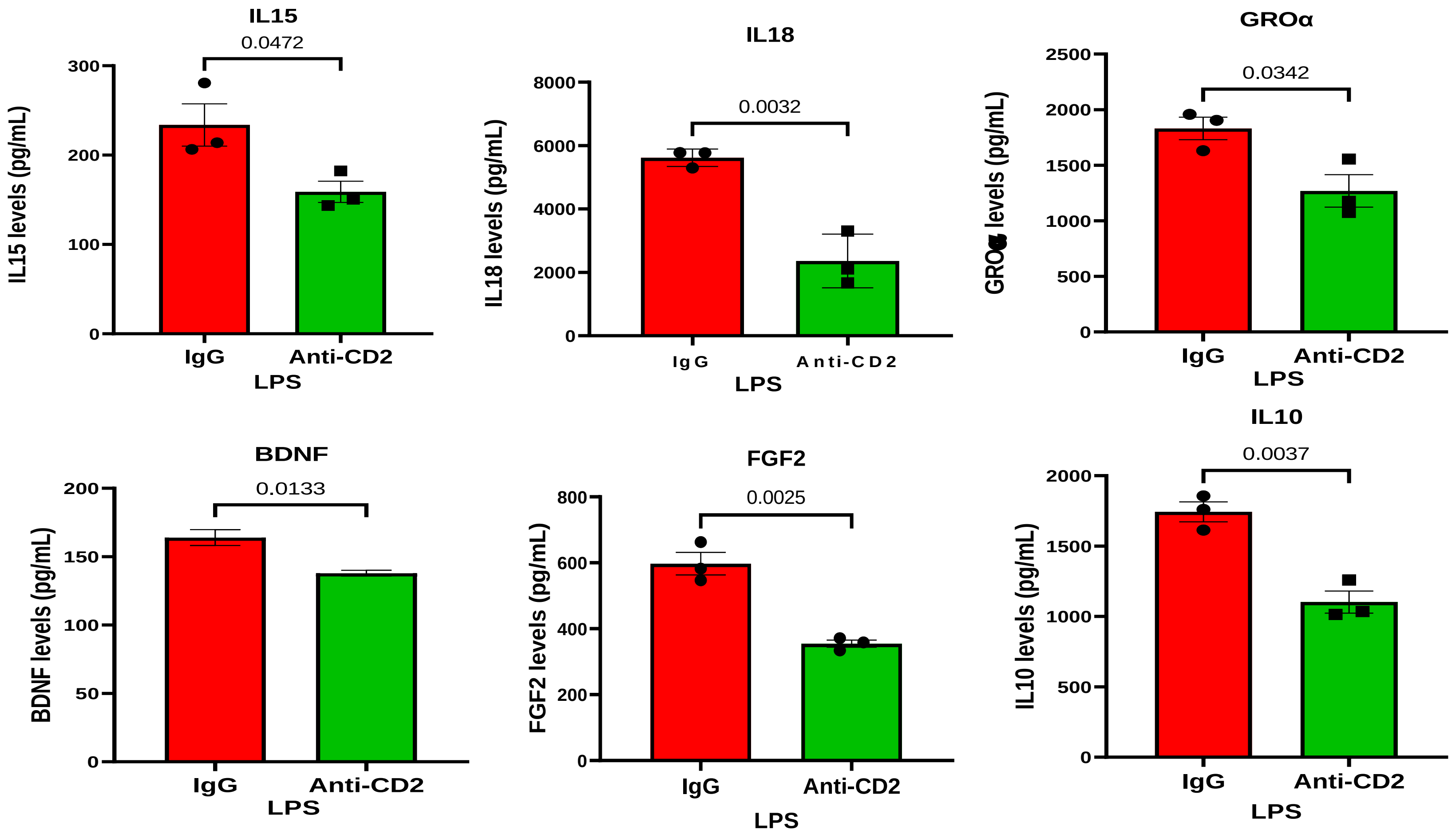


**Figure S10. Levels of cytokine, chemokines and growth factors in organotypic cultures of HS skin treated with LPS and anti-CD2 or IgG.**

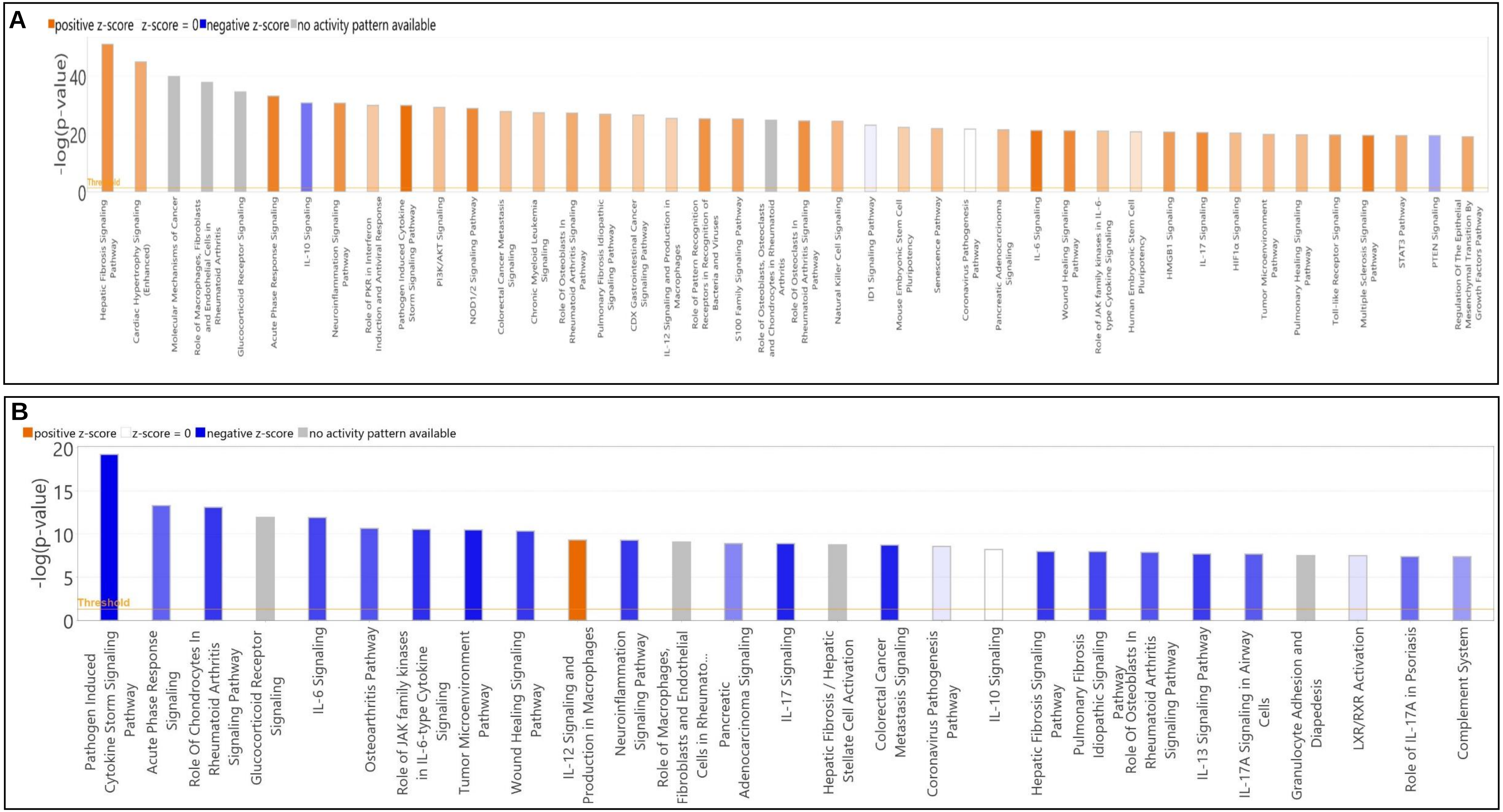


**Supplementary Figure S11. Activated or inhibited signaling/pathways of IPA analysis of HS vs normal (Figure 1D) and anti-CD2 or IgG (Figure 7).**

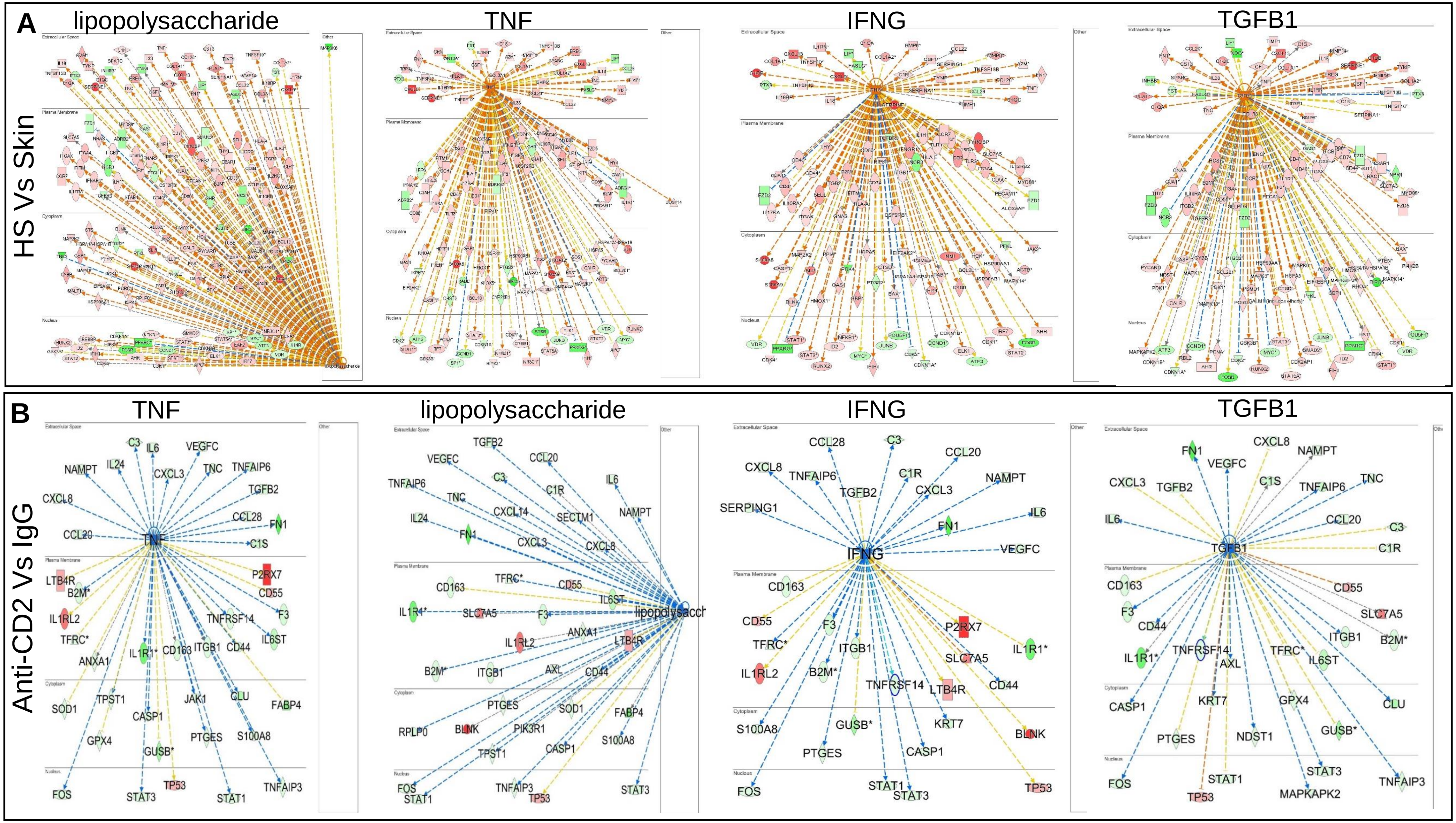


**Supplementary Figure S12. Upstream regulators as determined by IPA analysis. (A)** Upstream regulators of HS compared to normal OpenArray data from Figure 1D. The data shows top four significantly upregulated upstream regulators namely, lipopolysaccharide (P≤8.27E-81, z-score= 6.550), TNF (P≤6.13E-70, z-score= 4.994), IFNG (P≤3.96E-64, z-score=5.459) and TGFB1 (P≤1.12E-60, z-score=3.963) in lesional HS skin versus healthy controls. **(B)** Upstream regulators of anti-CD2 compared to IgG treated HS skin in organotypic cultures. The upstream regulators TNF (P≤1.70E-32, z-score= -3.721), lipopolysaccharide (P≤1.61E-28; z-score= -4.288), IFNG (P≤1.72E-27; z-score -2.993) and TGFB1 (P≤3.52E-26, z-score= -2.236) were significantly downregulated in HS skin treated with anti-CD2 relative to IgG.
